# Intrinsic mechanisms contributing to the biophysical signature of mouse gamma motoneurons

**DOI:** 10.1101/2025.06.29.661254

**Authors:** Simon A. Sharples, Struan J. Nisbet, Matthew J. Broadhead, Dennis Bo Jensen, Francesca L. Sorrell, Claire F. Meehan, Gareth B. Miles

## Abstract

Precise motor control relies on continuous sensory feedback from muscles, a process in which gamma motoneurons play a key role. These specialized spinal neurons innervate intrafusal muscle fibres, modulating their sensitivity to stretch and maintaining proprioceptive signalling during movement. Gamma motoneurons are characterized by a distinct biophysical profile, including low recruitment thresholds and high firing rates that enable rapid activation of intrafusal fibres at contraction onset. Despite their importance, the intrinsic mechanisms that underlie these properties remain poorly understood. In this study, we analysed published and unpublished data to identify a population of low-threshold, high-gain motoneurons with features consistent with gamma motoneurons, emerging during the third postnatal week in mice. Their low recruitment threshold was linked to lower membrane capacitance, higher input resistance, a more hyperpolarized activation of persistent inward currents (PICs), and a narrower axon initial segment. In contrast, higher firing rates were associated not with PIC amplitude but with shorter action potential durations and smaller medium afterhyperpolarizations. Notably, 92% of putative gamma motoneurons exhibited a sodium pump-mediated ultra-slow afterhyperpolarization (usAHP), which was absent in slow alpha motoneurons. This difference could not be attributed to h-current activity or expression of the alpha 3 subunit of the sodium-potassium ATPase. These findings reveal key intrinsic properties that support the unique excitability of gamma motoneurons, offering new insight into their contribution to motor control. This work provides a foundation for future studies into their development, regulation, and involvement in neuromuscular disorders.

**Key Points:**

● A distinct cluster of motoneurons with low recruitment current and high firing gain, characteristic of gamma motoneurons, emerges in the third week of postnatal development.
● Gamma motoneurons have a low recruitment current due to lower capacitance, higher input resistance, and a more hyperpolarized activation voltage for persistent inward currents.
● Their high firing rates are not driven by differences in persistent inward current amplitude but are instead attributed to shorter duration action potentials and smaller amplitude medium afterhyperpolarizations.
● A narrower axon initial segment in gamma motoneurons may contribute to their increased excitability compared to alpha motoneurons.
● Gamma motoneurons present with a higher prevalence of ultra slow afterhyperpolarization than slow alpha motoneurons that cannot be accounted for by differences in h-current or expression of alpha 3 subunits of the sodium potassium ATPase pump.

## Introduction

Motoneurons form the final common pathway by which signals from the central nervous system reach muscles (Eccles, 1957). As such, they are critical effectors of a wide range of motor behaviours. Two key mechanisms to grade the intensity of muscle contraction and movement vigour include the progressive recruitment of additional motoneurons and/or increasing their firing rates (Harrison, 1983; Kernell *et al*., 1999). Gamma motoneurons are a specialized subtype of motoneuron that innervate intrafusal fibres that are innervated by muscle spindles and form key proprioceptive sensory organs that detect muscle length. These motoneurons contribute little to the force generated during muscle contraction but have the ability to grade movement gain indirectly by adjusting muscle spindle sensitivity, which can in turn increase the excitability of alpha motoneurons via the monosynaptic reflex pathway. Gamma motoneurons possess a distinct biophysical profile that enables their earlier recruitment and higher firing rates compared to alpha motoneurons (Kemm & Westbury, 1978; Westbury, 1981; Khan *et al*., 2022). This signature is thought to promote rapid fusion of intrafusal muscle fibres, preserving spindle sensitivity during muscle contraction (Kemm & Westbury, 1978; Westbury, 1981). However, compared to alpha motoneurons, we have a limited understanding of the properties that underlie their early recruitment and high firing gain. This knowledge gap is partly due to their small size, making them a challenge to study using electrophysiological approaches in vivo, and the limited utility of in vitro models, which have typically been limited to early postnatal stages when spinal circuits are relatively immature. However, recent advances in approaches to identify gamma motoneurons (Manuel & Zytnicki, 2019; Wilkinson, 2021; Kang *et al*., 2024) and the development of in vitro spinal cord preparations from older animals (Hochman, 2011; Mitra & Brownstone, 2012; Özyurt *et al*., 2022) has opened new possibilities to study gamma motoneuron function.

Classically, gamma motoneurons have been identified as small cholinergic cells in the ventrolateral spinal cord that do not express NeuN or Hb9 and are devoid of cholinergic C bouton and Vglut1 synapses (Friese *et al*., 2009; Shneider *et al*., 2009; Manuel & Zytnicki, 2019; Wilkinson, 2021; Kang *et al*., 2024). Recent advances in single cell RNA sequencing approaches have provided insight into molecular markers of gamma motoneurons (Blum *et al*., 2021; Alkaslasi *et al*., 2021; Patel *et al*., 2022), with previous studies identifying Err2/3, Wnt7a, the GDNF receptor (GFR1α), and the alpha-3 subunit of the sodium-potassium ATPase pump found to be key molecular markers to identify gamma motoneurons (Friese *et al*., 2009; Shneider *et al*., 2009; Ashrafi *et al*., 2012; Edwards *et al*., 2013; Khan *et al*., 2022). Although the genetic identity of gamma motoneurons is established during embryonic development (Ashrafi *et al*., 2012) and present at birth (Shneider *et al*., 2009), it is unclear when gamma motoneurons become biophysically distinct from alpha motoneurons during development. This is important because identifying when this biophysical signature emerges during development is a pre-requisite to designing studies aimed at determining underlying mechanisms governing their function, neuromodulatory control, and contributions to motor dysfunction in disease. Given that this signature is particularly important for the dynamic control of complex movement (Khan *et al*., 2022), we hypothesized that gamma motoneurons become functionally distinct from alpha motoneurons at the start of the third postnatal week, a developmental stage when key molecular markers become confined to gamma motoneurons (Khan *et al*., 2022), and complex locomotor behaviours begin to emerge (Altman & Sudarshan, 1975; Khan *et al*., 2022).

To test this hypothesis, we reanalysed published data (Smith & Brownstone, 2020; Sharples & Miles, 2021; Sharples *et al*., 2023) and present new data to identify when gamma motoneurons become functionally distinct from alpha motoneurons during postnatal development. Consistent with our hypothesis, we identified a cluster of motoneurons exhibiting the biophysical signature of gamma motoneurons, which emerges at the start of the third postnatal week. This signature is mediated by a combination of passive and active properties, facilitating their early recruitment and action potential characteristics contributing to a higher firing gain across the frequency-current range compared to alpha motoneurons. Together, these findings enhance our understanding of a poorly characterized subtype of spinal motoneuron that plays a crucial role in coordinating complex movements.

## Results

We analysed published data (n = 270; (Sharples & Miles, 2021; Sharples *et al*., 2023)) and performed new recordings (n = 29 MNs) of lumbar motoneurons studied across the first three weeks of postnatal development. All data were obtained using whole-cell patch clamp recordings of motoneurons in transverse slices of the lumbar spinal cord. A subset of motoneurons were identified through retrograde labelling with fluorogold injected intraperitoneally to label and record from smaller motoneurons (Miles *et al*., 2005; Sharples & Miles, 2021). In these preparations, two motoneuron subtypes can be identified based on the onset and pattern of repetitive firing induced by long (5 s) depolarizing current steps applied near rheobase (Figure 1A). One subtype, which correlates with fast type motoneurons, exhibits a delayed onset to repetitive firing, and an accelerating firing rate. The other subtype, which is believed to correlate with slow type motoneurons, exhibits an immediate onset to repetitive firing with stable or adapting firing rates. Whilst multiple research groups have validated this approach for separating fast (delayed firing) and slow (immediate firing) motoneurons (Leroy *et al*., 2014, 2015; Bhumbra & Beato, 2018; Bos *et al*., 2018; Sharples & Miles, 2021; Nascimento *et al*., 2024), it has been challenging to separate other motoneuron subtypes; such as fatigue-sensitive versus fatigue-resistant fast motoneurons that both show delayed firing, or slow versus gamma motoneurons which would both be expected to exhibit immediate firing.

**Figure 1:**
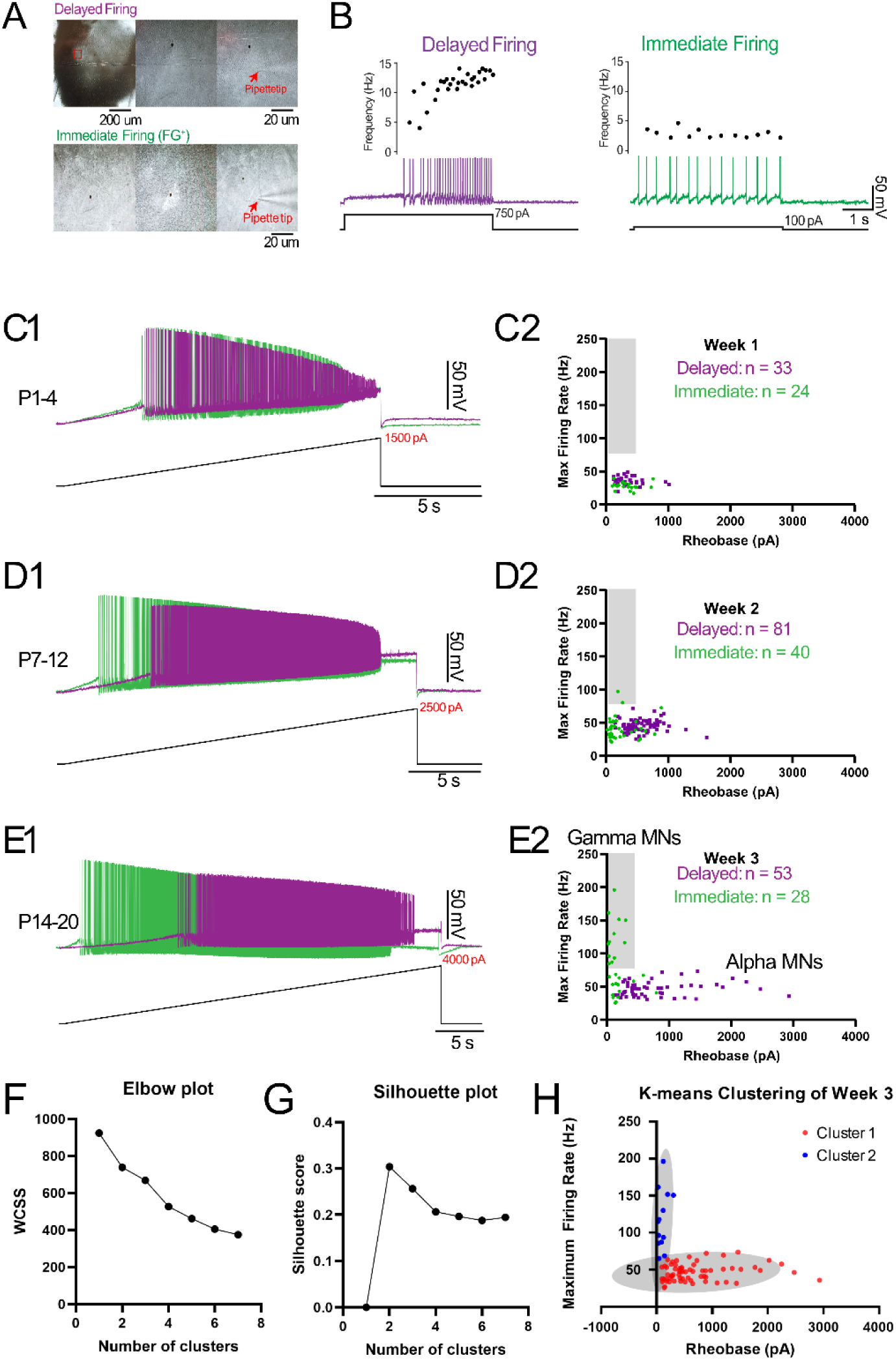
Emergence of a cluster of high excitability motoneurons during the third postnatal week that have a gamma-like biophysical signature. A) Whole cell patch-clamp recordings were obtained from motoneurons identified in the ventrolateral horn of the lumbar spinal cord under differential interference contrast with a subset retrogradely labelled with Fluorogold injected intraperitoneally. B) Motoneuron subtypes were identified based on delayed (purple) and immediate (green) onsets to repetitive firing during long (5 s) depolarizing current steps applied near rheobase current. C1-E1) The rheobase current of delayed and immediate firing motoneurons was assessed using slow (100 pA/s) depolarizing current ramps at weeks 1-3.C2-E2) Maximum firing rate plotted as a function of rheobase for delayed and immediate firing motoneurons during postnatal weeks 1-2. Number of motoneurons are highlighted in each graph. Light grey boxes in C2-E2 and F1-F3 delimit a zone occupied by putative gamma motoneurons defined by Khan et al., 2021. Pannels A-E are adapted from Sharples and Miles, 2021. (F) Elbow plot showing the total within-cluster sum of squares (WCSS) for increasing values of k (number of clusters). A clear inflection point at k = 2 indicates diminishing returns beyond two clusters, supporting a two-cluster solution.(G) Average silhouette width for clustering solutions with k = 2 to 6. The silhouette score peaked at k = 2 (0.31), suggesting this solution best represents the underlying structure of the data. (H) Scatter plot between rheobase and maximum firing rate with points colored according to k-means cluster assignment (Cluster 1: blue; Cluster 2: red; grey ellipses: 90% confidence intervals). The clustering reveals a distinct separation, with one cluster predominantly composed of immediate firing motoneurons characterized by low rheobase and high maximal firing rate, consistent with putative gamma motoneuron identity.

Therefore, in order to separate slow alpha motoneurons from gamma motoneurons, we first took a similar approach to Khan et al., (2022) and plotted rheobase against maximum firing rate, which we assessed during slow depolarizing current ramps (100 pA/s). This analysis allowed us to identify a cluster of immediate firing motoneurons with a low rheobase and high firing gain that might represent the gamma motoneuron population (Figure 1 C1-E1). Visual inspection of these data across the first three postnatal weeks revealed a cluster of immediate firing motoneurons with a low rheobase and a maximum firing rate greater than 80 Hz. This cluster emerged after P14 (Figure 1 C2-E2), suggesting that putative gamma motoneurons might become functionally diversified from slow alpha motoneurons at the start of the third postnatal week.

Having visually identified a subset of immediate firing motoneurons during the third postnatal week with biophysical properties consistent with gamma motoneurons, we next applied k-means clustering to statistically validate this classification. The analysis included 12 electrophysiological variables representing both passive and active membrane properties, recorded from 83 motoneurons (53 delayed firing, 30 immediate firing; Age: Age P14-20). The optimal number of clusters (k) was determined using an elbow plot, which showed a clear inflection at k = 2 (Figure 1F), and by evaluating silhouette scores, which peaked at 0.31 for k = 2 (Figure 1G). Together, these metrics supported a two-cluster solution as the most appropriate representation of the underlying structure in the data. Consistent with our initial observations, one of the clusters primarily contained immediate firing motoneurons characterized by a low rheobase and high maximal firing rate (Figure 1H)—properties that align with the known electrophysiological signature of gamma motoneurons. This clustering result provides statistical support for the presence of a physiologically distinct subset of gamma motoneurons within the immediate firing population.

We further validated this finding by analysing an open-access dataset published by Smith and Brownstone (2020), who studied maturation of firing properties in cervical and lumbar motoneurons (n = 138), identified using Hb9::eGFP mice, during a similar period of postnatal development. Given that gamma motoneurons do not express green fluorescent protein in Hb9::eGPF mice, we hypothesised that there should be an absence of putative gamma motoneurons in their recordings during the third postnatal week. Consistent with this hypothesis, and in support of our proposed identification of functional gamma motoneurons, analyses of data from Smith and Brownstone (2020) failed to reveal any motoneurons with a low rheobase and maximal firing rate above 80 Hz at any stage of postnatal development (Supplementary Figure 1 A - C).

Having identified a cluster of immediate firing motoneurons, that resemble gamma motoneurons during the third week, using both visual and unsupervised approaches, we next set out to determine key properties that might contribute to the lower rheobase and higher firing rates of putative gamma motoneurons (n = 18) compared to putative slow alpha motoneurons (n = 21; Figure 2A), which are summarized in Table 3. In line with what would be expected, putative gamma motoneurons had a significantly lower rheobase (Figure 2B)^a^, were recruited over a narrower current range (gMN: 278 pA sMN: 755 pA) and had a steeper recruitment gain compared to putative slow motoneurons. Putative gamma motoneurons also had a lower capacitance^b^ and higher input resistance^c^ compared to putative slow alpha motoneurons (Figure 2C & D), suggesting that differences in passive properties may contribute to the lower rheobase of gamma motoneurons. Further analysis of firing rates during depolarizing current ramps (Figure 2E&F) demonstrated that putative gamma motoneurons have a higher firing gain across the entire frequency-current range, supported by higher minimum^d^ and maximum^e^ firing rates and steeper frequency-current (f-I) slopes^f^ compared to slow alpha motoneurons (Figure 2G-I).

**Figure 2:**
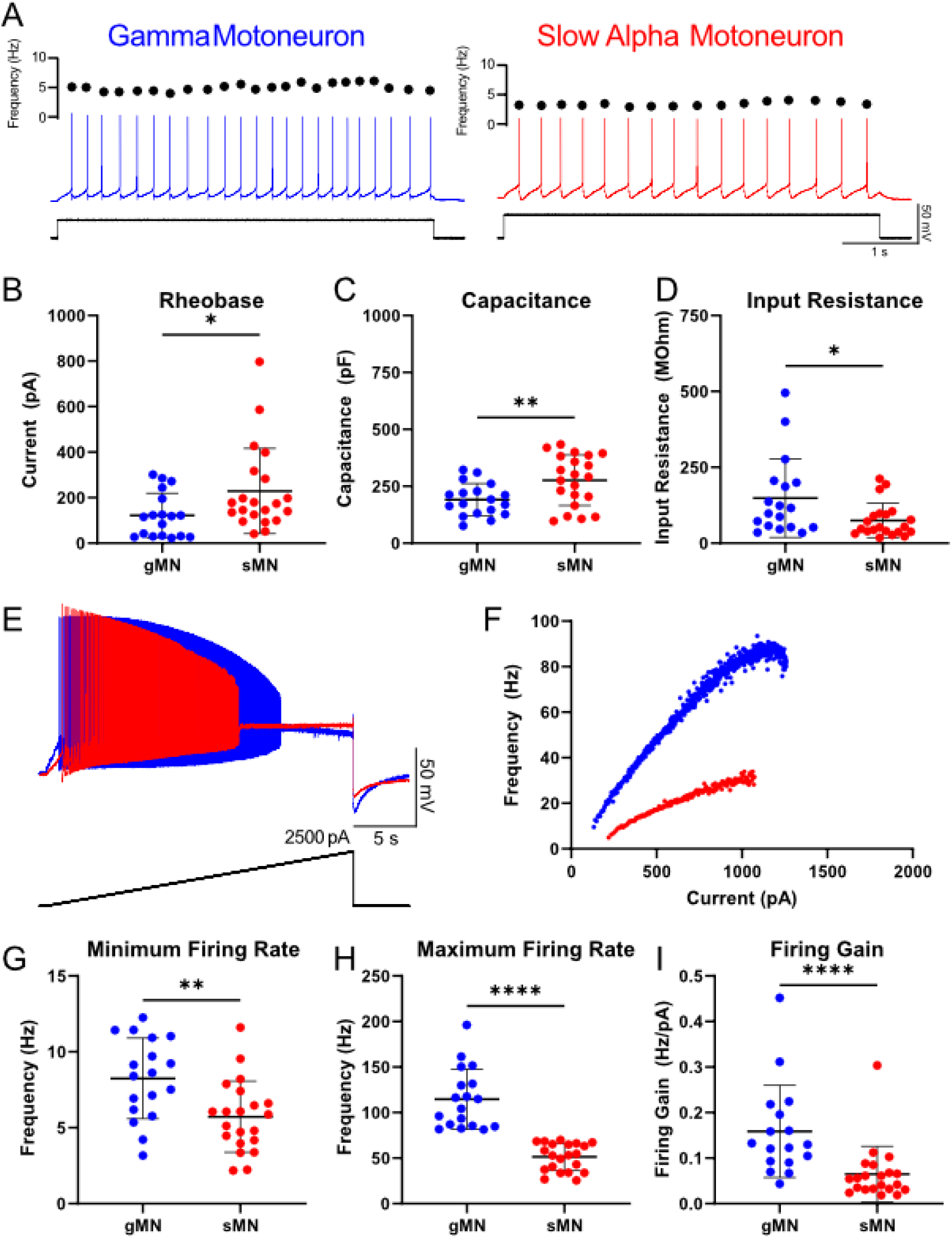
Intrinsic properties of immediate firing motoneurons composed of putative gamma and slow alpha motoneurons. A) Representative traces of immediate firing motoneurons from the putative gamma cluster (gMN - blue; n = 18) and slow alpha motoneurons (sMN - red; 21) during long depolarizing current steps applied near rheobase obtained from mice age during the third postnatal week. Putative gamma motoneurons have a lower rheobase (B), lower whole cell capacitance (C), and higher input resistance (D) compared to slow motoneurons. E) representative traces of a putative gamma motoneuron (blue) and slow alpha motoneuron (red) during slow depolarizing current ramps. F) Frequency current plot for representative gamma and slow alpha motoneurons. The minimum firing rate, maximum firing rate, and firing gain were higher in putative gamma motoneurons compared to slow alpha motoneurons. Individual data points are displayed, black bars represent mean ± SD. Statistical analysis was conducted using an unpaired t-test or an equivalent non-parametric test. Asterisks denote significant differences -p<0.05, **p<0.01, ***p<0.001, ****p<0.0001.

Motoneuron firing rates are influenced by persistent inward currents (PICs) (Schwindt & Crill, 1977; Lee & Heckman, 1998; Li & Bennett, 2003), which are largely conducted by Nav1.6/1.1 channels located on the axon initial segment (Brocard *et al*., 2016; Drouillas *et al*., 2023) and CaV1.3 channels on the distal dendrites (Jiang *et al*., 1999; Carlin *et al*., 2000*a*, 2000*b*; Heckmann *et al*., 2005; Elbasiouny *et al*., 2005, 2006). We recently demonstrated that NaV1.6 channels regulate motoneuron recruitment by controlling the activation voltage of PICs (Li *et al*., 2004; Sharples & Miles, 2021). We therefore hypothesized that higher firing rates and lower rheobase in putative gamma motoneurons could be supported by larger PICs that are activated at a more hyperpolarized voltage compared to slow alpha motoneurons. To test this hypothesis, we first estimated the impact of PIC on motoneuron firing using a triangular depolarizing current ramp and measured recruitment-derecruitment hysteresis (Bennett *et al*., 2001; Durand *et al*., 2015; Sharples & Miles, 2021) by calculating the difference in current at firing offset and onset on descending and ascending limbs of the ramp (Delta I) (Figure 3A). Although there was no significant difference in recruitment-derecruitment hysteresis^g^, qualitatively, putative gamma motoneurons (n = 12) always had a Delta I value above zero (Offset I > Onset Current) whereas putative slow alpha motoneurons (n = 15) exhibited a range of both positive and negative Delta I values (Figure 3B). Firing rate hysteresis can also be studied by comparing firing rate during ascending and descending limbs of the current ramp to identify different firing profiles that are indicative of non-linearities in motoneuron firing (Figure 3C). Further examination of non-linearities in firing hysteresis demonstrated that the firing rate of putative gamma motoneurons was often lower on the descending limb of the ramp compared to the ascending limb of the ramp, which is indicative of adaptive firing. In contrast, putative slow alpha motoneurons were more likely to have firing rates that were higher on the descending limb of the ramp compared to the ascending limb of the ramp, which is indicative of sustained firing and larger PIC, although there were also cells that produced adaptive firing (Figure 3D). These data suggest, in contrast to our hypothesis, that gamma motoneurons might have smaller PICs than slow alpha motoneurons.

**Figure 3:**
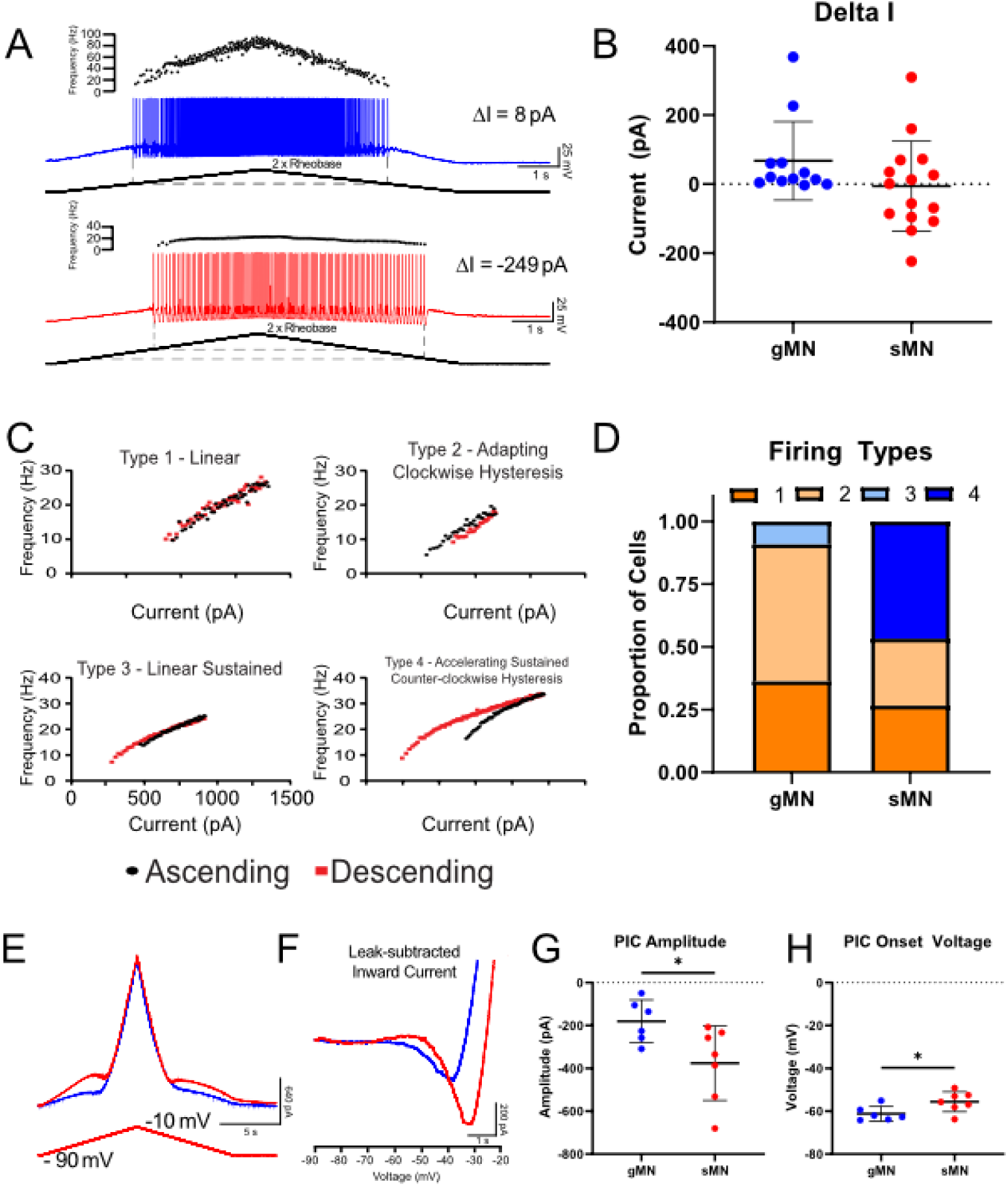
Smaller persistent inward current and adaptive firing in gamma motoneurons. A) PIC gives rise to non-linearities in motoneuron firing that can be detected during triangular current ramps where larger PIC allows motoneurons to fire at a lower current on the descending limb of the ramp compared to the ascending limb of the ramp (Delta I). B) Delta I did not differ between putative gamma and slow alpha motoneurons. C) Motoneurons demonstrate 4 firing types when comparing firing rates on ascending (black dots) and descending (red dots) limbs of triangular current ramps. These include type 1: linear, type 2: adapting, type 3: linear sustained, and type 4: accelerating sustained. D) Gamma motoneurons (n = 12) present with a higher proportion of type 1 and with 50% of slow alpha motoneurons (n = 15) demonstrating type 4 firing. E) Representative trace of a persistent inward current (PIC) measured from putative gamma motoneuron (gMN - blue) and slow alpha motoneuron (sMN - red) in voltage clamp using a depolarizing voltage ramp from a holding potential of -90 mV (black trace). F) Representative leak-subtracted traces of PICs. G) PIC was smaller and activated at more hyperpolarized voltages (H) in gamma compared to slow alpha motoneurons. Individual data points are displayed; black bars represent mean ± SD. Statistical analysis was conducted using an unpaired t-test or an equivalent non-parametric test. Asterisks denote significant differences p<0.05. Pannel C is adapted from Sharples and Miles, 2021.

To confirm this possibility, we next measured PICs more directly from putative gamma (n = 6) and slow alpha (n=7) motoneurons in voltage clamp using slow depolarizing voltage ramps (Figure 3E, F) (Quinlan *et al*., 2011; Steele *et al*., 2020; Reedich *et al*., 2023). In support of estimates of PIC in current clamp, but in contrast to our initial hypothesis, we found that putative gamma motoneurons do indeed have smaller PICs compared to putative slow alpha motoneurons^h^; however, there was no difference in maximal PIC density when these currents were scaled to capacitance (gMN = 1.0 ±0.6; sMN: 1.1 ± 0.45 pA/pF)^i^. Interestingly, the activation voltage of PICs was more hyperpolarized in putative gamma compared to putative slow alpha motoneurons (Figure 3G-H)^j^. Further analysis revealed that the PIC activation voltage was below the resting potential of most (4/6) putative gamma motoneurons (gMN RMP: -59.4 ± 6.2 mV) whereas PICs were activated above resting potential (sMN RMP: -65.5 ± 8.5 mV) in 6/7 putative slow alpha motoneurons. Together, these results suggest that a resting PIC may support a more depolarized resting potential^k^ and contribute to a lower rheobase in gamma motoneurons compared to slow alpha motoneurons. However, in contrast to our hypothesis, PICs do not seem to play a role in promoting high firing rates in gamma motoneurons.

Properties of individual action potentials and the medium afterhyperpolarization are well known to contribute to the firing rate of motoneurons across the frequency-current range (Eccles *et al*., 1957; Harrison, 1983; Kernell *et al*., 1999). We first examined whether differences in individual action potentials could account for higher firing rates in putative gamma motoneurons compared to putative slow alpha motoneurons across the frequency-current range. We found that putative gamma motoneurons had a lower rise time (Figure 4A1, A2)^l^ and narrower action potential half-width (Figure 4A3)^m^ compared to putative slow alpha motoneurons. We next examined properties of the medium afterhyperpolarization (mAHP), which is also known to limit firing rates (Kernell, 1965; Bakels & Kernell, 1993). We found that putative gamma motoneurons had a lower amplitude mAHP than putative slow alpha motoneurons (Figure 4B1-B2)^n^; however, there was no difference in mAHP half-width (Figure 5B3)°. Together these results suggest that differences in neuronal properties that influence action potential shape and mAHP amplitude may contribute to higher firing rates reached by gamma motoneurons across the entire frequency-current range.

**Figure 4:**
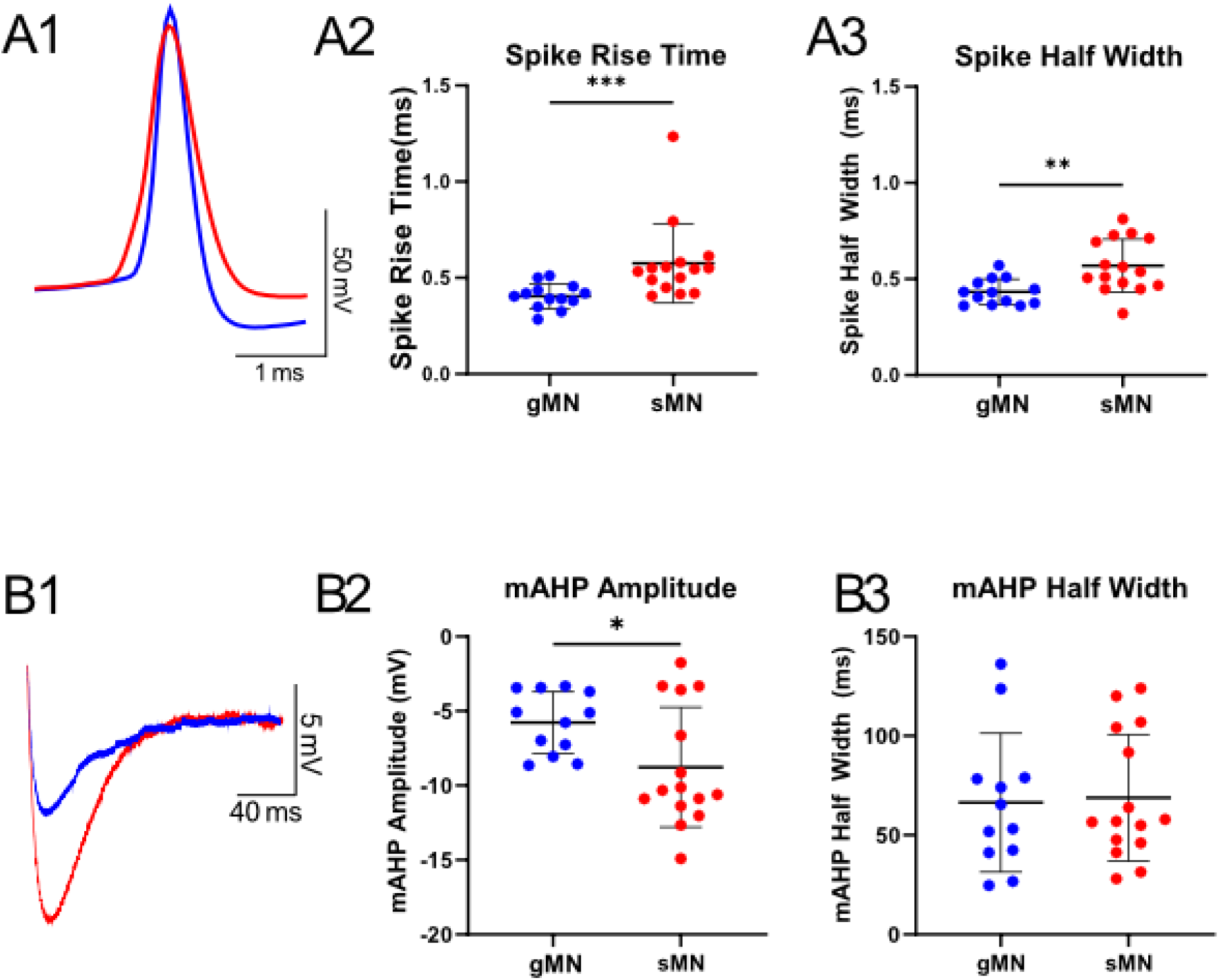
Action potentials are shorter and medium afterhyperpolarization (mAHP) smaller in putative gamma compared to slow alpha motoneurons. A1) Representative trace of a single action potential from putative gamma (gMN - Blue; n = 13) and slow alpha motoneuron (sMN - red; n = 15) elicited in response to short (10 ms) depolarizing current steps applied at an intensity of 1.5 times rheobase. Gamma motoneurons have a shorter action potential rise time (A2) and half width (A3). Representative trace of the medium after hyperpolarization (mAHP) from putative gamma and alpha slow motoneurons. The mAHP was significantly larger in slow motoneurons (B2) but did not differ in mAHP half width (B2). Individual data points are displayed; black bars represent mean ± SD. Statistical analysis was conducted using an unpaired t-test or an equivalent non-parametric test. Asterisks denote significant differences -p<0.05, **p<0.01, ***p<0.001.

**Figure 5:**
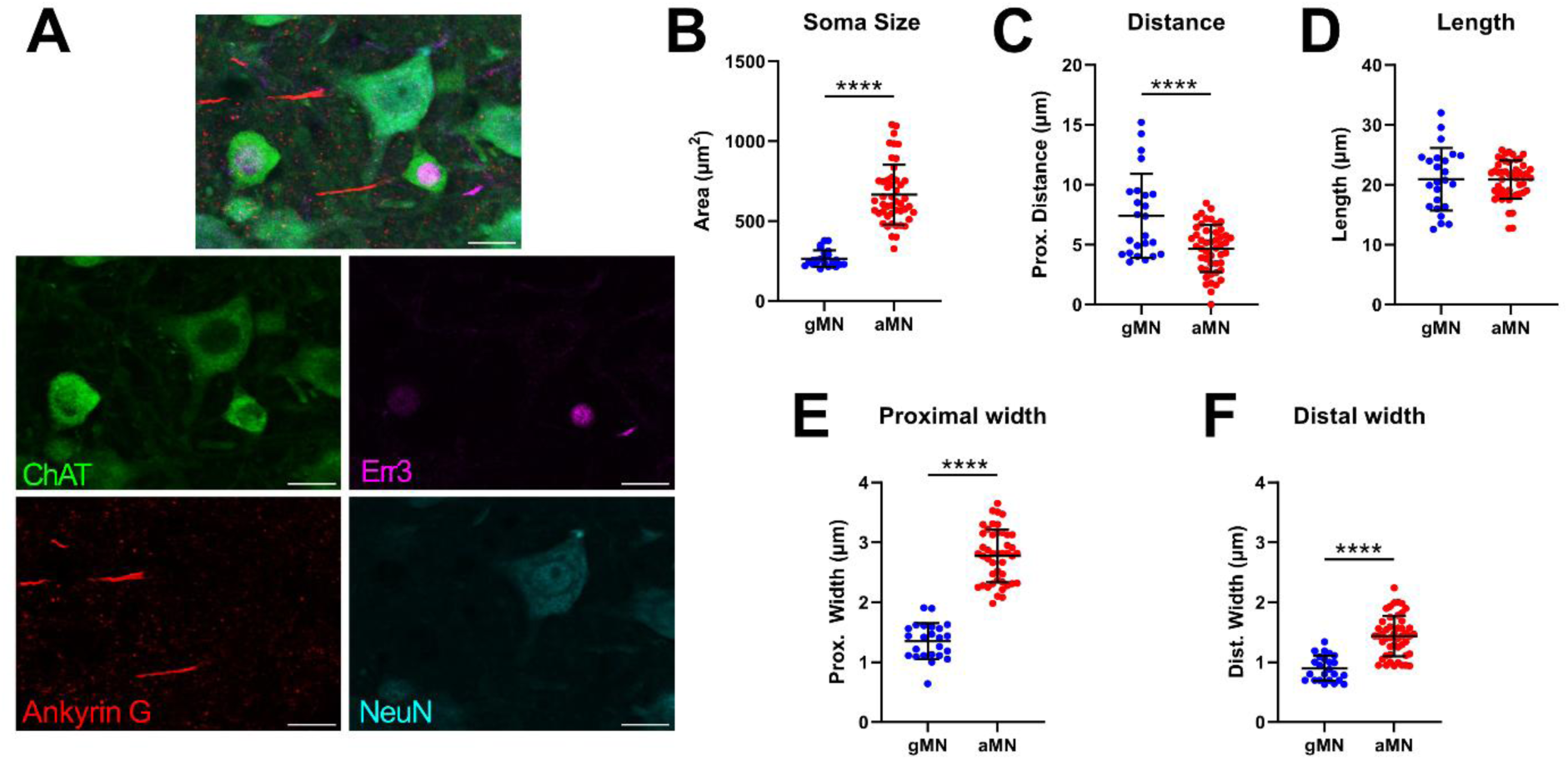
Gamma motoneurons have a narrower axon initial segment (AIS) than alpha motoneurons. A) Immunohistochemical labelling for Choline Acetyl Transferase (ChAT; green), Estrogen-related receptor 3 (Err3; magenta), Ankyrin G (red) and NeuN (cyan) with scale bars = 20 µm. Gamma motoneurons were identified based on co-expression of ChAT and Err3 and were devoid of NeuN labelling, whereas alpha motoneurons were identified based on co-expression of ChAT and NeuN and were devoid of Err3 labelling. B) Gamma motoneurons (gMN; blue dots; n = 24 MNs, 6 animals) had a smaller soma area compared to alpha motoneurons (aMN; red dots; n = 48 MNs, 6 animals). C) The proximal border of the AIS, labelled with Ankyrin G, of gMNs was further from the soma but was the same length compared to aMNs (D). The AIS of gMNs was narrower at both proximal (E) and distal borders (F) compared to aMNs. Individual datapoints represent MNs with mean±SD displayed. Statistical analysis was conducted using an unpaired t-test. Asterisks denote significant differences - **** p<0.0001.

In our next experiment, we aimed to examine whether differences in geometrical properties could also contribute to the increased excitability of gamma motoneurons compared to alpha motoneurons. Previous studies have shown that gamma motoneurons have a smaller soma size and a less complex dendritic arbor, which may contribute to their heightened excitability. However, another key structure, the axon initial segment (AIS), where voltage-gated ion channels necessary for action potential generation are clustered (Verneuil *et al*., 2020; Rotterman *et al*., 2021), has not been characterized in gamma motoneurons. To address this gap, we tested the hypothesis that differences in AIS geometry could contribute to the increased excitability of gamma motoneurons. Motoneurons were identified using Choline Acetyltransferase (ChAT) labeling, while AISs were labeled with antibodies against the Ankyrin G protein, which spans the entire initial segment (Jensen *et al*., 2020; Jørgensen *et al*., 2021; Djukic *et al*., 2025). Alpha and gamma motoneurons were distinguished based on the expression of the transcription factor Err3, which is highly expressed in the nuclei of gamma motoneurons but not in alpha motoneurons, and NeuN labeling, which is abundant in the nuclei and cytoplasm of alpha motoneurons but absent in gamma motoneurons (Figure 5A) (Friese *et al*., 2009; Khan *et al*., 2022).

Consistent with previous reports (Friese *et al*., 2009; Khan *et al*., 2022) and our electrophysiological measurements of capacitance and input resistance (Figure 2), the somata of gamma motoneurons (Err3+, NeuN-, ChAT+; n = 24 cells, 6 animals) had a smaller area than those of alpha motoneurons (Err3-, NeuN+, ChAT+; n = 48 cells, 6 animals) (Figure 5B)^p^. However, contrary to our hypothesis, Ankyrin G labeling revealed that the proximal border of the AIS was significantly farther from the soma in gamma motoneurons (Figure 5C)^q^, while the AIS length remained unchanged compared to alpha motoneurons (Figure 5D)^r^. Interestingly, we also found that the AIS was narrower at both its proximal (Figure 5E)^s^ and distal (Figure 5F)^t^ borders in gamma motoneurons, a feature that may support their increased excitability. This result was the same whether statistical analyses were conducted using individual motoneurons or average measurements within animals^u-w^. Together, these results suggest that specific geometric features of the AIS may contribute to the enhanced excitability of gamma motoneurons compared to alpha motoneurons.

In our final set of experiments, we set out to validate the alpha-3 subunit of the sodium potassium ATPase pump as a marker of gamma motoneurons. Previous work in adult mice has demonstrated that gamma motoneurons can be identified by the expression of the alpha-3 subunit of the sodium potassium ATPase pump (Edwards *et al*., 2013). This isoform of the sodium potassium pump has a lower affinity for sodium than the tonically active alpha-1 subunit. As a result, the activity of sodium potassium pumps composed of alpha-3 subunits are increased when the intracellular sodium level is elevated, such as occurs following intense and prolonged neuronal firing. Due to the asymmetric movement of charge, it is believed that alpha-3-expressing sodium pumps produce a prolonged hyperpolarization of the membrane potential, known as the ultra-slow afterhyperpolarization (usAHP), that does not change membrane input resistance and recovers over a period of up to a minute following sustained firing (Pulver & Griffith, 2010; Zhang *et al*., 2015; Picton *et al*., 2017; Akkuratov *et al*., 2025). However, previous studies have found that motoneurons present with a diverse array of post-discharge activities, with a subset of motoneurons displaying the pump mediated usAHP, whilst other motoneurons present with a slow AHP (sAHPs) or afterdepolarizations (ADPs) (Bouhadfane *et al*., 2013; Picton *et al*., 2017; Bos *et al*., 2021; Drouillas *et al*., 2023; Harris-Warrick *et al*., 2024; Akkuratov *et al*., 2025). We therefore hypothesized that gamma motoneurons would be enriched in pump mediated usAHPs, with alpha motoneurons presenting with more sAHPs and ADPs.

In line with previous reports, we found that usAHPs in immediate firing motoneurons increased in magnitude with action potential frequency (Figure 6A), did not alter the input resistance (Figure 6B), and could be blocked by low doses of ouabain (Figure 6C1-4, 1 uM, n = 6)^x-z^. Having confirmed that the usAHP in immediate firing motoneurons is pump mediated, we next set out to determine whether differences in usAHP magnitude between slow and putative gamma motoneurons would be in line with what would be expected based on the expression of the Alpha-3 subunits of the sodium pump in gamma motoneurons (Edwards *et al*., 2013). We therefore compared the post-discharge activities of putative gamma motoneurons to putative slow, alpha motoneurons. In line with what would be expected based on previously published work (Edwards *et al*., 2013), 92% of the putative gamma motoneurons (n = 12) presented with a usAHP following a 10 second depolarizing current step applied at an intensity of 2 times rheobase. In contrast, no slow alpha motoneurons (n = 12) presented with a usAHP but instead produced neutral responses (50%), or in fewer cases presented with slow afterhyperpolarizations (25%) or slow afterdepolarizations (25%). Overall, the post-discharge activity of gamma motoneurons had a larger area^aa^, were longer in duration^bb^, and larger in amplitude^cc^ than the slow alpha motoneurons (Figure 6C2-C4).

**Figure 6:**
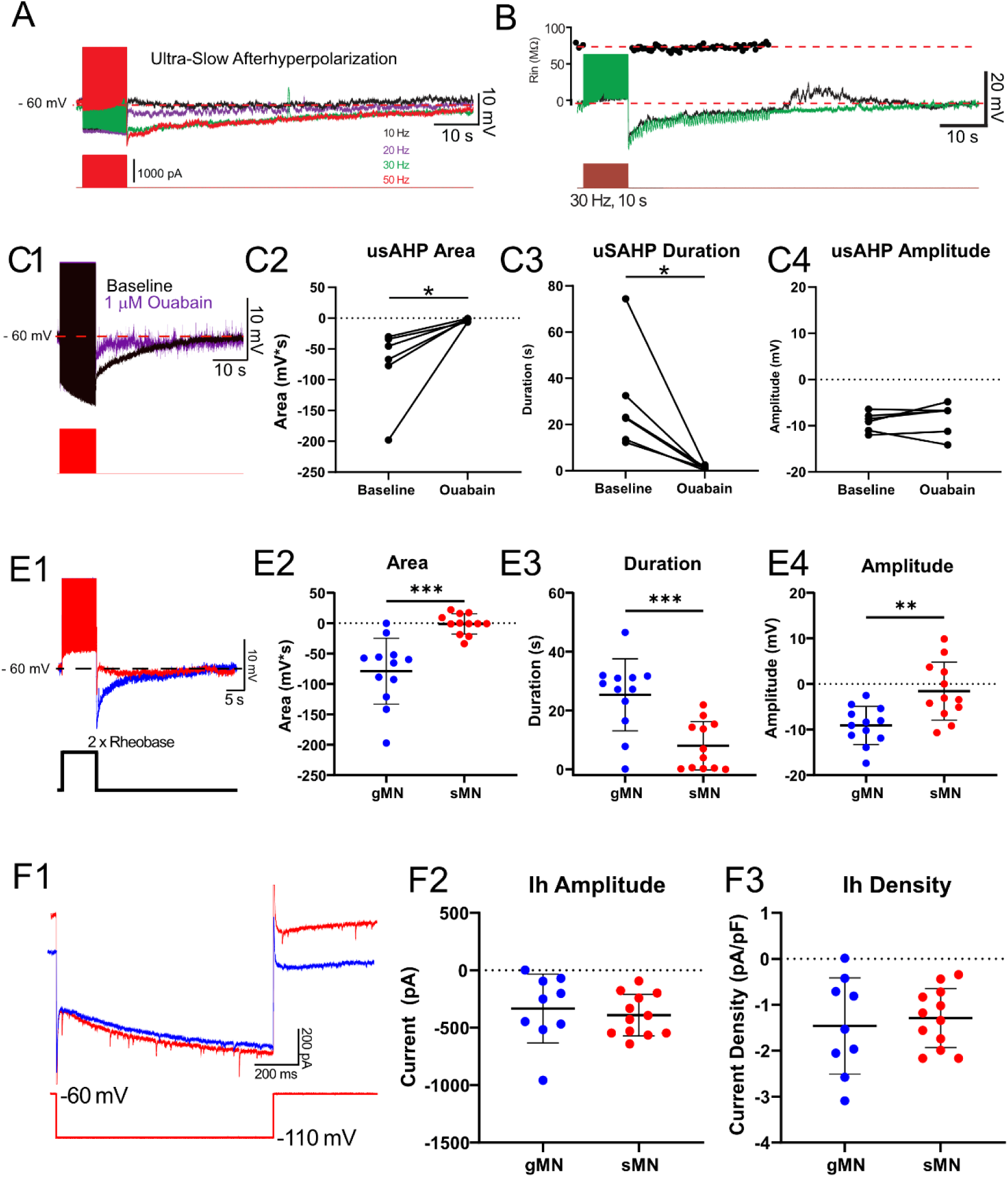
The sodium potassium ATPase-pump mediated ultra-slow AHP (usAHP) is larger in putative gamma compared to slow alpha motoneurons. A) usAHP increases in amplitude and duration in response to increased stimulation frequency. B) Input resistance remains constant during the usAHP, as measured in response to hyperpolarizing current steps (green).C1) Representative trace of the usAHP in response to 10 second train (30 Hz) before (black) and after a low dose (1 uM; n = 6) of ouabain (purple). Ouabain reduces usAHP area (C2), duration (C3) but does not affect peak usAHP amplitude (C4). E1) Representative trace of a usAHP in response to a 10s depolarizing current step applied at an intensity of 2 times rheobase. Gamma motoneurons displayed characteristics of an ultra-slow afterhyperpolarization, which had a larger and more negative area (E2), longer duration (E3), larger amplitude (E4). F1) Representative trace of a hyperpolarization-activated inward current (Ih) measured in voltage clamp during hyperpolarizing voltage steps. F2) Ih amplitude and density (F3) did not differ between putative gamma (n = 9) and slow alpha (n = 12) motoneurons. Individual data points are displayed, black bars represent mean ± SD. Statistical analysis was conducted using an unpaired t-test or an equivalent non-parametric test. Asterisks denote significant differences -p<0.05, **p<0.01, ***p<0.001.

We next considered the possibility that differences in the post-discharge activities between gamma and slow alpha motoneurons may be mediated by ion channels that oppose the inhibitory actions of the sodium-potassium ATPase pump. For example, in *Xenopus* spinal interneurons, the usAHP is opposed and masked by a hyperpolarization activated inward current (Ih) (Picton *et al*., 2018) that is active at resting potential and could therefore account for differences in the prevalence or magnitude of the post-discharge activities that we observe. However, in contrast to this hypothesis, we did not find evidence of a resting Ih current in either cell type and there was no difference in maximal Ih amplitude^dd^ or current density^ee^ (Figure 6D1-D3), in putative gamma (n = 9) and slow alpha (n = 12) motoneurons.

We next sought to validate the α3 subunit of the sodium-potassium ATPase as a bona fide marker of gamma motoneurons and to determine whether differences in α3 expression between gamma and small alpha motoneurons could explain the variation in usAHPs observed in our electrophysiological recordings during the third postnatal week. To address this, motoneurons were labeled in P18 mice by intraperitoneal injection of the retrograde tracer Fluorogold, and the α3 subunit was visualized via immunohistochemistry using specific antibodies. Alpha and gamma motoneurons were distinguished based on two established criteria: differential Fluorogold uptake—reported to be higher in gamma motoneurons than in alpha motoneurons (Khan *et al*., 2022)—and NeuN immunoreactivity, which is robust in the nuclei and cytoplasm of alpha motoneurons but absent in gamma motoneurons (Figure 7A, 7B). To focus on relevant populations, we selectively analyzed motoneurons with soma areas below 300 μm² (Friese *et al*., 2009; Harris-Warrick *et al*., 2024), allowing us to assess α3 subunit expression in gamma motoneurons (high Fluorogold, NeuN-negative) and putative slow alpha motoneurons (low Fluorogold, NeuN-positive). Consistent with prior findings (Khan *et al*., 2022), we identified small NeuN-negative neurons with high levels of Fluorogold uptake, supporting their classification as gamma motoneurons. The soma size of these cells was comparable to the gamma motoneurons identified using Err3 labeling in our earlier experiments, providing convergent validation of this classification approach. Contrary to our hypothesis, however, immunolabeling revealed that both gamma motoneurons (high FG, NeuN−) and slow alpha motoneurons (low FG, NeuN+) exhibited membrane localization of the α3 subunit (Figure 7B), with no significant difference in labeling intensity between the two cell types (Figure 7C)^ff.^ These findings suggest that the enhanced prevalence of usAHPs in gamma motoneurons during the third postnatal week is unlikely to be driven by differential expression of the α3 subunit.

**Figure 7:**
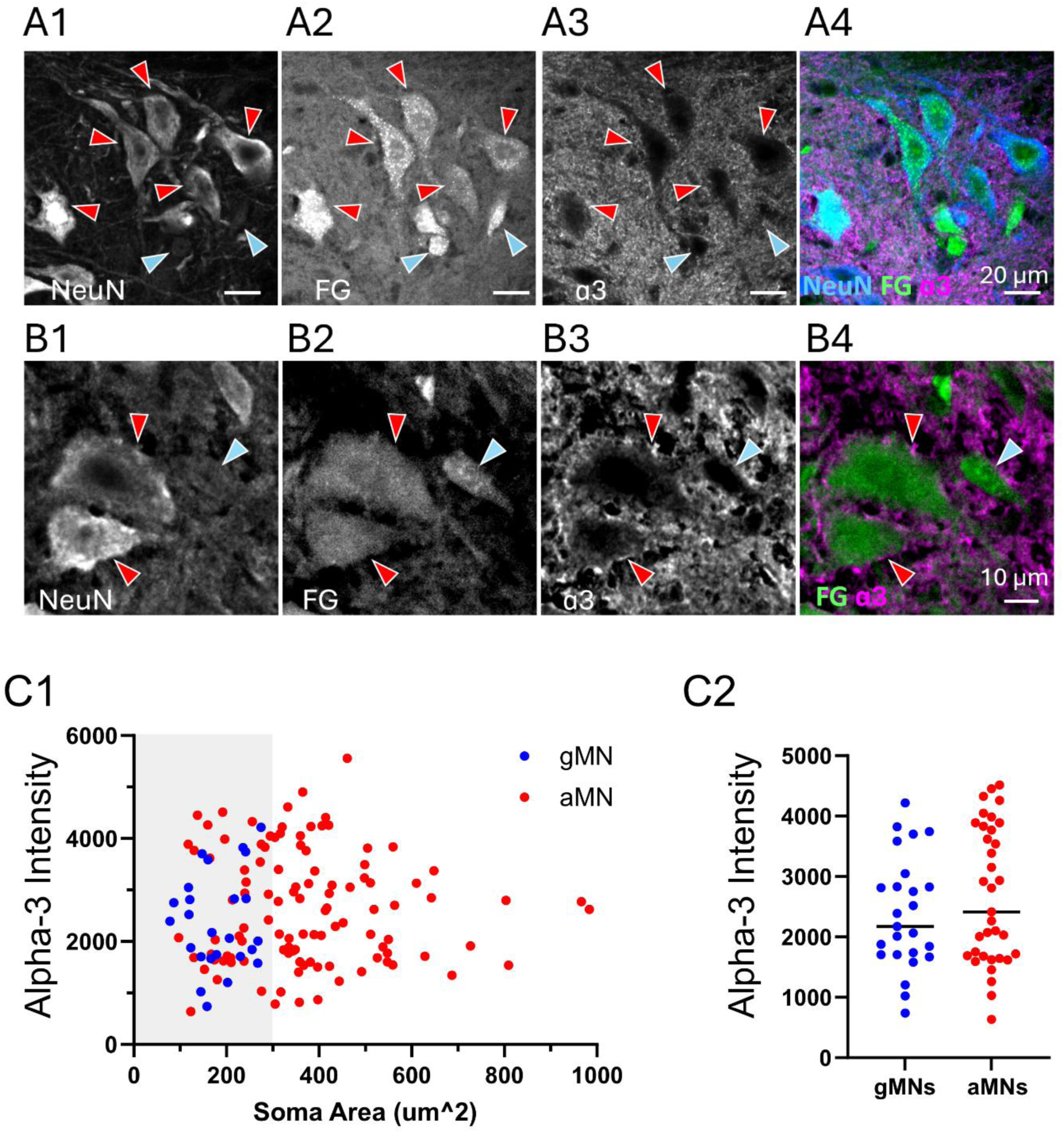
Both gamma and slow alpha motoneurons express the alpha 3 subunit of the sodium potassium ATPase pump at the third postnatal week. A) Image of lateral motor pool of lumbar spinal cord tissue from a P18 mouse labelled for NeuN (A1), Fluorogold (“FG”, A2), and the ɑ3 subunit of the sodium potassium ATPase pump (A3). Composite image shown in A4. B) High magnification image of alpha and gamma motoneurons labelled for NeuN (B1), Fluorogold (“FG”, B2), and ɑ3 subunit (B3). Composite image shown in B4. Gamma MNs (blue arrows) were identified as bright FG +ve cells with low NeuN labelling and a small size (<300 µm2). Alpha MNs (red arrows) were identified as larger cells, modest FG intensity, and higher intensity of NeuN labelling. C1) Scatter plot of the cell size (µm^2) and membrane-associated ɑ3 subunit labelling in putative gamma (blue), alpha (red) MNs from P18 animals (n=133 MNs, 4 animals). C2) All MNs <300 µm2 were analysed and separated based on whether they were classed as Gamma MNs (n=25) or alpha MNs (n=35) from 4 mice at P18. C2) Membrane associated ɑ3 subunit expression was not significantly different between Gamma and alpha MNs. F) Data in panels C2 were analysed using a Mann Whitney U.

**Supplementary Figure 1:**
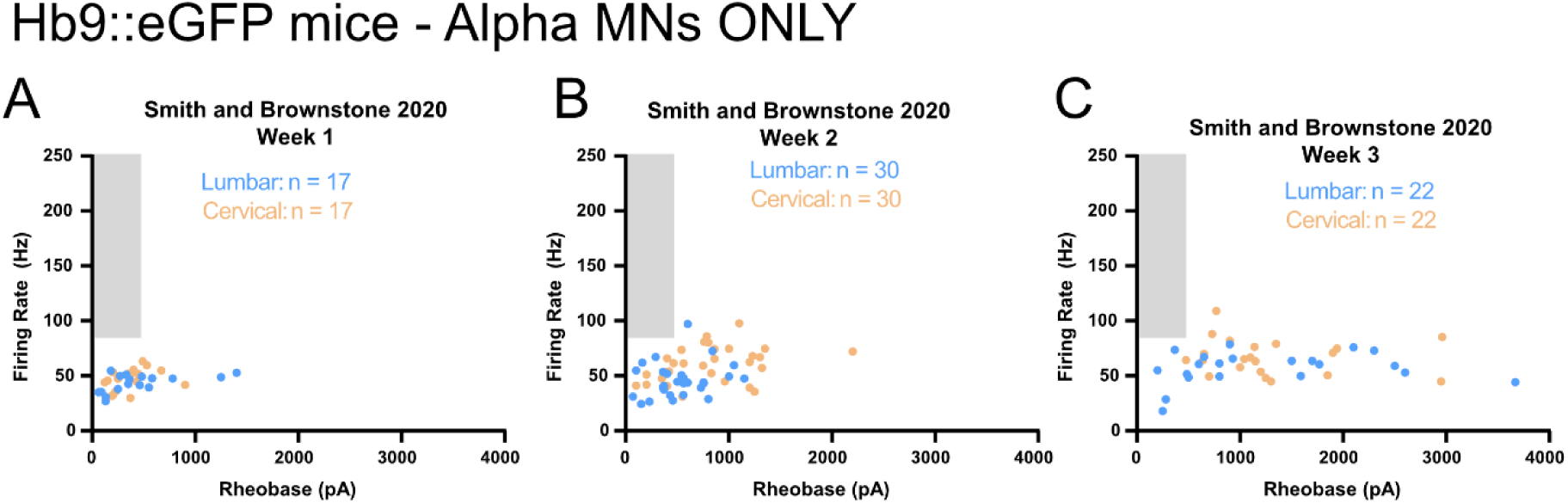
A-C) Maximum firing rate assessed during depolarizing current steps and plotted as a function of rheobase for Hb9::GFP+ motoneurons in lumbar (blue) and cervical (orange) spinal segments during postnatal weeks 1-3 studied by Smith and Brownstone (2020).

## Discussion

Gamma motoneurons are a functionally distinct subtype of motoneurons that remain poorly understood compared to alpha motoneurons (Manuel & Zytnicki, 2019; Wilkinson, 2021; Kang *et al*., 2024). Despite a known biophysical signature of gamma motoneurons (Kemm & Westbury, 1978; Westbury, 1981; Khan *et al*., 2022), the intrinsic properties that support their early recruitment and high maximal firing rates are poorly understood. This biophysical signature of gamma motoneurons is particularly important for maintaining spindle sensitivity during movement and for supporting the generation of complex motor behaviors (Khan *et al*., 2022). In this study, we identified intrinsic mechanisms that may contribute to early recruitment and higher firing rates of gamma compared to alpha motoneurons. We leveraged datasets from both published and unpublished recordings collected from studies of motoneuron maturation across the first three weeks of postnatal development in mice (Smith & Brownstone, 2020; Sharples & Miles, 2021; Sharples *et al*., 2023). These datasets are particularly valuable due to the significant challenges in obtaining viable motoneuron recordings in vitro during postnatal stages when mice produce more mature movements and provide a unique opportunity to determine when gamma motoneurons become functionally distinct from alpha motoneurons. Through our visual and unbiased statistical analyses, we identified a cluster of motoneurons with a low rheobase and high firing rates that emerged during the third postnatal week, which matched the biophysical signature of gamma motoneurons (Kemm & Westbury, 1978; Westbury, 1981; Khan *et al*., 2022). We further corroborated this observation by analyzing published data from Smith and Brownstone, (2020), who studied maturation of motoneurons in Hb9::eGFP mice across a similar period of postnatal development. In line with what would be expected, given that gamma motoneurons do not express GFP in Hb9::eGFP mice (Shneider *et al*., 2009; Koronfel *et al*., 2021), we did not find any cells with a low rheobase and high firing rate at any stage of postnatal development, thus supporting our initial identification of putative gamma motoneurons. Although speculative, it is possible that the functional maturation of gamma motoneurons may contribute to the emergence of complex motor behaviours during the third week of postnatal development (Altman & Sudarshan, 1975). Having identified this cluster of putative gamma motoneurons, we further investigated key properties and currents that may contribute to this biophysical signature, which are discussed below.

### Recruitment

Alpha-gamma coactivation is a key mechanism for ensuring the appropriate activation of the fusimotor system during movement (Granit, 1975; Ellaway *et al*., 2015; Niyo *et al*., 2024). While the shared premotor inputs that may mediate this kind of connectivity are not yet defined, the lower rheobase of gamma motoneurons, compared to alpha motoneurons, may ensure their recruitment prior to alpha motoneurons, helping to maintain spindle sensitivity at the onset of muscle contraction. In line with Henneman’s Size Principle (Henneman, 1957), and previous work (Friese *et al*., 2009; Khan *et al*., 2022), we show that Err3+, NeuN-labelled gamma motoneurons are substantially smaller than Err3-, NeuN+ alpha motoneurons. This observation is further supported by our electrophysiological measurements demonstrating lower capacitance and higher input resistance of putative gamma motoneurons. However, in line with our measurements of whole cell capacitance, some anatomical studies have also shown considerable overlap in soma size between gamma motoneurons and the smallest alpha motoneurons (Shneider *et al*., 2009; Morita-Isogai *et al*., 2017; Khan *et al*., 2022), suggesting that additional properties may contribute to the early recruitment of gamma motoneurons. Our anatomical studies suggest that, in addition to a smaller soma size, gamma motoneurons also have a narrower axon initial segment, which may contribute to their earlier recruitment. As in our previous work comparing fast and slow motoneurons (Sharples & Miles, 2021), we also identified a role for active properties in facilitating the recruitment of putative gamma prior to alpha motoneurons. Notably, we observed that persistent inward currents (PICs) in putative gamma motoneurons are activated at more hyperpolarized voltages compared to putative slow alpha motoneurons. In fact, in 66% of putative gamma motoneurons studied, PICs were activated below the resting membrane potential, suggesting that a resting PIC may support a more depolarized resting potential. This observation is in line with electrophysiological studies in hypoglossal motoneurons, which demonstrate a hyperpolarization of the resting potential following application of the NaV channel blocker, Riluzole (Bellingham, 2013). While our recordings were made at a standardized membrane potential, the observed hyperpolarized PIC activation could suggest differences in resting potential that may contribute further to earlier recruitment in vivo. Together these results provide novel insights into mechanisms that support the early recruitment of gamma motoneurons.

### Firing Rate

Early recordings of nerve impulses from the ventral roots of anesthetized cats provided the first evidence that gamma motoneurons fire at a significantly higher rate than alpha motoneurons (Kemm & Westbury, 1978; Westbury, 1981). This elevated maximal firing rate is thought to facilitate the rapid fusion of muscle spindles upon recruitment, maintaining spindle sensitivity at the onset of muscle contraction. More recently, similar findings have been demonstrated in mouse and chick motoneurons studied *in vitro*, where Kv1.8 shaker channels have been shown to support higher maximal firing rates of gamma motoneurons (Khan *et al*., 2022). However, our results indicate that putative gamma motoneurons fire at a higher rate across the entire frequency-current (FI) range, likely due to a broader complement of voltage-gated ion channels that regulate their firing gain. For instance, the duration or amplitude of the medium afterhyperpolarization (mAHP) plays a key role in limiting firing rate in motoneurons and is also a key property that distinguishes different motoneuron types (Gardiner & Kernell, 1990; Bakels & Kernell, 1993; Deardorff *et al*., 2013; Rotterman *et al*., 2021). It has even been postulated that gamma motoneurons may lack a long-lasting mAHP (Gustafsson & Lipski, 1979). We found that the mAHP is indeed present in putative gamma motoneurons and is relatively smaller compared to that in slow alpha motoneurons. This reduced mAHP may be due to differences in the expression of channels underlying the mAHP. Although debated (Kissane *et al*., 2022), it is generally believed that the larger mAHP in slow compared to fast motoneurons is mediated by differences in SK2 and SK3 subunit expression (Deardorff *et al*., 2013). Therefore, it is also possible that a smaller mAHP in gamma compared to slow motoneurons may also be mediated by differences in SK2 and SK3 subunit expression.

We also considered additional properties that may contribute to the higher firing rates observed in gamma motoneurons across the FI range. Initially, we hypothesized that larger PICs might support the elevated firing rates of gamma motoneurons. However, contrary to this hypothesis, PICs were smaller in putative gamma motoneurons compared to putative slow alpha motoneurons. Instead, we found that putative gamma motoneurons exhibit narrower action potentials, which may support a higher firing rate at the upper end of the frequency-current range, where the mAHP contributes relatively less. In line with this notion, our previous work has reported that modulatory C-bouton synapses can increase maximal firing rate of motoneurons by reducing spike width (Miles *et al*., 2007; Nascimento *et al*., 2020; Eleftheriadis *et al*., 2023). Channels such as Kv1.8, which have been shown to be enriched in gamma motoneurons (Khan *et al*., 2022), may also contribute to this narrowing, though further work is needed to directly assess their involvement. We also hypothesized that differences in the properties of the axon initial segment (AIS) might contribute to higher firing gain. In theory, one might assume that a more distal axon initial segment would increase the electrical isolation of the initial segment and soma. Yet, modelling studies have suggested this is not so straightforward, with the functional consequence of a more distal local depending on the soma size (Gulledge & Bravo, 2016). These suggest that with small neurons, a more proximal initial segment may increase excitability and for larger cells such as alpha motoneurons, then a more distal location may be more advantageous. From this, one could conclude that the different location of the initial segments between alpha and gamma motoneurons might simply reflect an adaptation to the different soma size to maintain optimal excitability. Surprisingly, in vivo recordings from rat alpha motoneurons within a single motor pool with anatomical measurements of their axon initial segments observed that motoneurons with more proximal axon initial segments tended to have lower somatic current thresholds for activation (Rotterman *et al*., 2021). Although this sample was within the alpha motoneuron pool where other factors also contribute to the orderly recruitment of the differential functional classes of motoneurons (Sharples & Miles, 2021; Sharples *et al*., 2023). Therefore, establishing a causal relationship may be challenging.

One feature that can influence this is axon initial segment diameter. It has been shown that neurons with larger soma tend to also have larger diameter axon initial segments and modelling suggests that, if sodium channel density remains the same, then this may be a strategy to maintain a uniform action potential threshold across different neuron classes (Goethals & Brette, 2020). Thus, our more distal, but smaller diameter axon initial segment may be crucial in coordinating its recruitment relative to the alpha motoneurons to ensure their optimal co-activation.

More recent modeling studies have shown that the effects of initial segment location may also depend on the axonal resistance between the soma and the initial segment (Fékété *et al*., 2021). This was demonstrated that decreasing the diameter of the axon between the soma and the initial segment may lower the voltage threshold, a measure of excitability which the author argues reflects differences in axon initial segment better than measures of rheobase (Fekete et al 2021). Thus, our observation that the diameter difference at the proximal axon initial segment diameter was greater between alpha and gamma motoneurons than that measured at the distal end of the initial segment suggests that this may contribute to the increased excitability of the gamma motoneurons. Ultimately, the effect will be influenced by the relative distribution of the different ion channels found at the axon initial segment. Therefore, our findings lay a crucial foundation for further investigation into other ion channels expressed at the AIS that may contribute to the distinctive biophysical signature of gamma motoneurons.

### Novel markers for gamma motoneurons

A fundamental challenge in studying gamma motoneurons is the lack of effective strategies for their identification and manipulation (Wilkinson, 2021; Rotterman *et al*., 2021; Kang *et al*., 2024). One difficulty is that the expression of many of the molecular markers change throughout postnatal development as motoneurons become functionally (Nakanishi & Whelan, 2010; Sharples & Miles, 2021) and transcriptionally diversified (Patel *et al*., 2022). For example, ERR2 (Errβ) and ERR3 (Errγ) are broadly expressed across multiple motoneuron subtypes at birth but become restricted to gamma motoneurons after the first few weeks of postnatal development (Friese *et al*., 2009; Shneider *et al*., 2009; Khan *et al*., 2022). Alternatively, gamma motoneurons can be identified by their small size or the absence of NeuN expression, although NeuN is weakly expressed in gamma motoneurons at birth (Shneider *et al*., 2009). In our labelling studies, we identified gamma motoneurons during the third postnatal week based on their small size, selective expression of Err3, absence of NeuN, and retrograde labelling with fluorogold, which has been shown previously to accumulate at higher levels in gamma motoneurons compared to alpha motoneurons (Khan *et al*., 2022). Despite these challenges, there are a few molecular markers present at birth that may facilitate the study of gamma motoneurons. For example, gamma motoneurons can be identified based on the expression of 5HT1D receptors (Enjin *et al*., 2012), GDNF (Gfrα1) receptors (Shneider *et al*., 2009), Wnt7a (Ashrafi *et al*., 2012). However, the developmental profile of many of these markers has not been explored in detail.

Our findings demonstrate a pronounced enrichment of ultra-slow afterhyperpolarizations (usAHPs) which may be mediated by the α3 subunit of the Na⁺/K⁺-ATPase in putative gamma motoneurons (Akkuratov *et al*., 2025). This result aligns with previous anatomical evidence of α3 expression in this cell type (Edwards *et al*., 2013) and introduces a novel biophysical signature that may aid in the functional identification of gamma motoneurons. The presence of the sodium pump-dependent usAHP in gamma motoneurons likely holds physiological relevance, serving as a form of inhibitory regulation for a cell type characterized by high intrinsic excitability, while maintaining sensitivity to synaptic inputs. Interestingly, despite the functional correlation between the usAHP and the α3 isoform, our data did not reveal a differential expression pattern of the α3 subunit that could account for the enrichment of usAHPs in putative gamma motoneurons. This discrepancy may reflect developmental differences, as our study focused on juvenile mice (P18), whereas prior reports examined adult animals (6–8 weeks of age). This raises the possibility that the molecular profile of gamma motoneurons undergoes further refinement beyond the third postnatal week. However, within the developmental window studied here, variation in α3 subunit expression does not explain the observed biophysical distinction between putative gamma and slow alpha motoneurons. Additional mechanisms may also contribute to the expression or masking of usAHPs. Studies in *Xenopus* spinal neurons have shown that a hyperpolarization-activated inward current (Ih) that is active at resting potential can obscure the usAHP, unmasking it upon HCN channel blockade (Picton *et al*., 2018). While our previous work has identified a subset of lumbar motoneurons that exhibit sodium pump-mediated usAHPs (Picton *et al*., 2017; Akkuratov *et al*., 2025), we have also demonstrated that fast alpha motoneurons develop a prominent resting H-current during the third postnatal week, which delays their recruitment (Sharples & Miles, 2021). In contrast, we found no evidence for a similar resting H-current in either putative gamma or slow alpha motoneurons at this age. This suggests that other, yet unidentified conductances may act to suppress or counterbalance the sodium pump’s influence specifically in slow alpha motoneurons. Furthermore, the sodium pump-mediated usAHPs observed in putative spinal gamma motoneurons in our study appear to differ from those described in gamma motoneurons innervating cranial muscles. For instance, gamma motoneurons projecting to the masseter muscle in the brainstem exhibit a post-pulse afterdepolarization (ADP) rather than a hyperpolarization. This ADP is mediated by a calcium-activated sodium current (Nishimura *et al*., 2018; Kang *et al*., 2024), resembling the ADPs observed in lumbar fast alpha motoneurons (Bouhadfane *et al*., 2013; Bos *et al*., 2021; Harris-Warrick *et al*., 2024). These contrasting findings suggest that gamma motoneurons may exhibit region- and muscle-specific electrophysiological diversity, likely reflecting the specialized mechanical demands and control strategies of different motor pools. Finally, recent studies have implicated mutations in the ATP1A3 gene, which encodes the α3 subunit of the sodium-potassium pump, in the pathogenesis of rapid-onset dystonia-parkinsonism (RDP)—a movement disorder characterized by abrupt motor hyperactivity (Akkuratov *et al*., 2025). Given our findings that gamma motoneurons are enriched in usAHPs, we speculate that sodium pump dysfunction could result in gamma motoneuron hyperexcitability, potentially contributing to the abnormal motor output observed in RDP. Future work should explore the role of gamma motoneuron physiology in the etiology of such motor syndromes.

## Conclusion

We leveraged existing datasets collected across multiple laboratories and new recordings to gain insight into the maturation of gamma motoneuron biophysical properties and to identify key features that contribute to their distinct biophysical signature. This work provides foundational information that will pave the way for future studies further investigating the mechanisms of gamma motoneuron properties, their neuromodulatory control, and their role in motor dysfunction following neural injury and disease. These studies will be further facilitated by recent advances in molecular markers that enable the selective identification, recording, and manipulation of gamma motoneurons (Blum *et al*., 2021; Alkaslasi *et al*., 2021; Patel *et al*., 2022; Karekal *et al*., 2025; Kussick *et al*., 2025).

## Methods

### Animals

This study included unpublished data in addition to reanalysis of experimental data summarised in (Smith & Brownstone, 2020; Sharples & Miles, 2021; Sharples *et al*., 2023). These published studies included experiments performed on tissue obtained from 133 (male: n = 68; and female: n = 65) wild type C57Bl/6J mice at postnatal days (P) 1-20 in Sharples and Miles, 2021, 10 mice (P14-18) from Sharples et al., 2023, and 54 Hb9::eFGP mice (P2-21) (B6.Cg-Tg(Hlxb9-GFP)1Tmj/K) from Smith and Brownstone, 2020. All procedures performed were conducted in accordance with the UK Animals (Scientific Procedures) Act 1986 and were approved by Animal Ethics Committees at the University of St Andrews and University of Copenhagen.

### Tissue preparation for in vitro spinal slice preparations

Animals were eviscerated and pinned ventral side up in a dissecting chamber lined with silicone elastomer (Sylguard) filled with carbogenated (95% oxygen, 5% carbon dioxide), ice-cold (1-2 degrees Celsius) potassium gluconate based dissecting/slicing aCSF (containing in mM: 130 K-gluconate, 15 KCl, 0.05 EGTA, 20 HEPES, 25 D-glucose, 3 kynurenic acid, 2 Na-pyruvate, 3 myo-inositol, 1 Na-L-ascorbate; pH 7.4, adjusted with NaOH; osmolarity approximately 345 mOsm). Spinal cords were exposed by performing a ventral laminectomy, cutting the ventral roots and gently lifting the spinal cord from the spinal column. Spinal cords were removed within 3 - 5 minutes following cervical dislocation. Spinal cords were secured directly to an agar block (3 % agar) with VetBond surgical glue (3M) and glued to the base of the slicing chamber with cyanoacrylate adhesive. The tissue was immersed in ice-cold dissecting/slicing aCSF bubbled with carbogen (95% O2, 5% CO2). Blocks of frozen slicing solution were also placed in the slicing chamber to keep the solution around 1-2 degrees Celsius. On average, the first slice was obtained within 10 minutes of decapitation which increased the likelihood of obtaining viable motoneurons in slices. 300 µm transverse slices were cut at a speed of 10 um/s on the vibratome (Leica VT1200) to minimise tissue compression during slicing. 3-4 slices were obtained from each animal. Slices were transferred to a recovery chamber filled with carbogenated pre-warmed (35 degrees Celsius) recovery aCSF (containing in mM: 119 NaCl, 1.9 KCl, 1.2 NaH2PO4, 10 MgSO4, 1 CaCl, 26 NaHCO3, 20 glucose, 1.5 kynurenic acid, 3% dextran) for thirty minutes after completion of the last slice which took 10 -15 minutes on average. Following recovery, slices were transferred to a chamber filled with carbogenated warm (35 degrees Celsius) recording aCSF (containing in mM: 127 NaCl, 3 KCl, 1.25 NaH2PO4, 1 MgCl, 2 CaCl2, 26 NaHCO3, 10 glucose) and allowed to equilibrate at room temperature (maintained at 23-25 degrees Celsius) for at least one hour before experiments were initiated.

### Whole-cell patch clamp electrophysiology

Slices were stabilized in a recording chamber with fine fibres secured to a platinum harp and visualized with a 40x objective with infrared illumination and differential interference contrast. A large proportion of the motoneurons studied were identified based on location in the ventrolateral region with somata greater than 20 µm. Recordings were obtained from a subset of motoneurons that had been retrogradely labelled with Fluorogold (Fluorochrome, Denver, CO). Fluorogold was dissolved in sterile saline solution and 0.04 mg/g injected intraperitoneally 24-48 hours prior to experiments (Miles *et al*., 2005; Sharples & Miles, 2021). In addition to recording from larger FG-positive cells, this approach allowed us to more confidently target smaller motoneurons. Motoneurons were visualized and whole cell recordings obtained under DIC illumination with pipettes (L: 100 mm, OD: 1.5 mm, ID: 0.84 mm; World Precision Instruments) pulled on a Flaming Brown micropipette puller (Sutter instruments P97) to a resistance of 2.5-3.5 MΩ. Pipettes were back-filled with intracellular solution (containing in mM: 140 KMeSO4, 10 NaCl, 1 CaCl2, 10 HEPES, 1 EGTA, 3 Mg-ATP and 0.4 GTP-Na2; pH 7.2-7.3, adjusted with KOH).

Signals were amplified and filtered (6 kHz low pass Bessel filter) with a Multiclamp 700 B amplifier, acquired at 20 kHz using a Digidata 1440A digitizer with pClamp Version 10.7 software (Molecular Devices) and stored on a computer for offline analysis.

### Identification of fast and slow motoneuron types

Motoneuron subtypes were identified using a protocol established by (Leroy *et al*., 2014), which differentiates motoneuron type based on the latency to the first spike when injecting a 5 second square depolarizing current near the threshold for repetitive firing. Using this approach we were able to identify 2 main firing profiles - a delayed repetitive firing profile with accelerating spike frequency, characteristic of fast-type motoneurons, and an immediate firing profile with little change in spike frequency, characteristic of slow-type motoneurons, which may also include gamma motoneurons (Figure 1; (Leroy *et al*., 2014)).

### K-means cluster analysis

To objectively validate the presence of a physiologically distinct subset of immediate firing motoneurons identified through visual inspection, we performed an unsupervised k-means clustering analysis using Graph Pad Pro Version 10.5 (Prism, San Diego, CA, USA). This analysis included a total of 83 motoneurons recorded during the third postnatal week (n = 53 delayed firing, n = 30 immediate firing). Twelve electrophysiological variables were selected to represent both passive and active membrane properties, including resting membrane potential, input resistance, membrane time constant, rheobase, spike threshold, spike amplitude, afterhyperpolarization amplitude and duration, firing latency, firing gain, maximal firing rate, and spike frequency adaptation index. Prior to clustering, all variables were standardized by z-transformation (mean = 0, standard deviation = 1) to ensure comparability across different measurement scales. K-means clustering was conducted using the Hartigan-Wong algorithm with Euclidean distance as the similarity metric. To ensure solution stability and avoid local minima, clustering was repeated with 100 random initial centroid initializations. The optimal number of clusters (k) was determined by visual inspection of an elbow plot showing the within-cluster sum of squares across increasing values of k, and by calculating the average silhouette width for each clustering solution. The elbow plot indicated a distinct inflection at k = 2, and the silhouette score was highest at k = 2 (average silhouette width = 0.31), supporting a two-cluster solution. Final cluster assignments were compared to initial firing type classifications to assess correspondence with visually identified immediate firing motoneurons.

### Acquisition and analysis of electrophysiology data

Data from whole cell patch clamp recordings were re-analysed from 408 motoneurons across the three studies (Smith & Brownstone, 2020; Sharples & Miles, 2021; Sharples *et al*., 2023). 259 lumbar motoneurons were studied from Sharples and Miles, 2021, 11 lumbar motoneurons from Sharples et al., 2023, and 138 GFP-expressing motoneurons at cervical (n = 70 MNs) or lumbar (n = 68 MNs) segments from Smith and Brownstone, 2020. Unpublished data from 29 lumbar motoneurons were analysed in the same manner as published data.

All motoneuron intrinsic properties were studied by applying a bias current to maintain the membrane potential at -60 mV. Values reported are not liquid junction potential corrected to facilitate comparisons with previously published data. Cells were excluded from analysis if access resistance was greater than 20 MΩ or changed by more than 5 MΩ over the duration of the recording, if resting membrane potential deviated by more than 5 mV over the duration of the recording period, or if spike amplitude measured from threshold (described below) was less than 60 mV.

Passive properties including capacitance, membrane time constant (tau), and passive input resistance (Ri) were measured during a hyperpolarizing current pulse that brought the membrane potential from - 60 to -70mV. Input resistance was measured from the initial voltage trough to minimize the impact of slower acting, active conductance (eg. Ih, sag). The time constant was measured as the time it took to reach 2/3 of the peak voltage change. Capacitance was calculated by dividing the time constant by the input resistance (C=T/R). Resting membrane potential was measured 10 minutes after obtaining a whole cell configuration and at the end of recordings from the MultiClamp Commander.

Rheobase and repetitive firing properties were assessed during slow (100 pA/s) depolarizing current ramps which allowed us to measure the recruitment current and voltage threshold of the first action potential. The voltage threshold of the first action potential was defined when the change in voltage reached 10 mV/ms. Firing rates were measured at rheobase and derecruitment (maximum firing rate) during slow depolarizing current ramps in Sharples and Miles, (2021) and Sharples et al., (2023), or during 1 second square depolarizing current steps in Smith and Brownstone, (2020).

Single spikes and the medium afterhyperpolarization (mAHP) were elicited using a 10 ms square, depolarising current pulse applied at an intensity 25% above rheobase current. Spike threshold was determined as the potential at which the derivative (dV/dt) increased above 10 mV/ms. Spike amplitude, rise time, and half width were measured from this point. Medium afterhyperpolarization (mAHP) properties (amplitude and half width) were measured from baseline (−60 mV). Single spike and mAHP properties were measured using event detection features in Clampfit Version 11 (Molecular Devices). The ultra-slow afterhyperpolarization (usAHP) was elicited and studied by injecting 10 depolarizing current steps at an intensity of 2 times rheobase or in response to a 10 second train of depolarizing current pulses. Post discharge responses were classified as an afterdepolarization (ADP), neutral, slow afterhyperpolarization (sAHP), or usAHP as in (Akkuratov *et al*., 2025).

Non-linearities in motoneuron firing as a result of persistent inward currents (PICs) were studied in current clamp mode using the injection of a triangular, depolarizing current ramp with 5 second rise and fall times (Li & Bennett, 2003; Li *et al*., 2004; Durand *et al*., 2015; Sharples & Miles, 2021). Depolarizing current ramps were set to a peak current of 2 times repetitive firing threshold current (determined with a 100pA/s depolarizing current ramp initiated from -60 mV). The influence of persistent inward currents on motoneuron excitability was estimated by calculating the difference (delta I) between the current at firing onset on the ascending component of the ramp and the current at derecruitment on the descending component of the ramp. A negative delta I is suggestive of a PIC. Firing hysteresis was also assessed on the ascending and descending components of the ramp. We subdivided cells into 1 of 4 types based on the firing hysteresis on ascending and descending portions of the ramp (Li *et al*., 2004; Durand *et al*., 2015; Sharples & Miles, 2021) (Type 1: Linear, Type 2: Adapting, Type 3: Sustained, Type 4: Counter clockwise).

PICs were measured in voltage clamp by injecting a slow depolarizing voltage ramp (10 mV/s from - 90 to -10 mV) over 8 seconds. PIC onset voltage, peak current amplitude, and peak current density were measured from post-hoc leak-subtracted traces as in (Quinlan *et al*., 2011; Steele *et al*., 2020; Verneuil *et al*., 2020; Reedich *et al*., 2023). Ih was measured in voltage clamp during 1 second hyperpolarizing voltage steps from - 60 mV to - 110 mV, in 10 mV increments. Ih was measured as the difference between the initial and steady state current.

### Immunohistochemistry and Microscopy

Two separate immunohistochemistry (IHC) protocols were used to examine α3 subunit expression and axon initial segment (AIS) morphology in mouse spinal motoneurons.

### Tissue Preparation and Perfusion

For experiments examining α3 subunit expression, four unsexed P9 and four female P17 C57Bl/6J mice received a single intraperitoneal injection of Fluorogold (FG; Fluorochrome) diluted in H₂O at a dose of 0.04 mg/g body weight. After 24 hours, mice were anesthetized with pentobarbital (30 mg/kg, Dolethal) and transcardially perfused with phosphate-buffered saline (PBS) followed by 4% paraformaldehyde (PFA). Spinal cords were post-fixed for 6 hours, transferred to 30% sucrose in PBS for 2–3 nights, embedded in OCT compound, and stored at –80°C. Thirty-micrometer (30 µm) thick cryosections were obtained using a NX50 cryostat (Epredia) and mounted on adhesive histology slides.

For AIS morphology studies, six P20 C57Bl/6J mice (3 male, 3 female) were deeply anesthetized and perfused with PBS followed by 4% PFA in pH 7.2 phosphate buffer. Spinal cords were removed, post-fixed in the same fixative for 2 hours, and cryoprotected overnight in 30% sucrose. Fifty-micrometer (50 µm) thick horizontal sections of the lumbar spinal cord were cut on a HM450 freezing microtome (Thermo Scientific) for free-floating immunohistochemistry.

### Immunohistochemistry Procedures

For α3 expression experiments, tissue sections were thawed to room temperature and washed in PBS before incubation in blocking solution containing 3% bovine serum albumin (BSA) and 0.2% Triton X-100 for 1.5 hours. Sections were then incubated for 2 nights at 4°C with primary antibodies (Table 1) diluted in PBS with 1.5% BSA and 0.1% Triton X-100. After washing in PBS, secondary antibodies (Table 1) were applied for 2 hours at room temperature in PBS with 0.1% Triton X-100. Sections were washed, dried, and coverslipped using ProLong Glass Antifade Mounting Medium (Thermo Fisher).

**Table 1:** Table of Antibodies

| Primary Antibody Target | Species | Company | Code | Dilution | Secondary Antibody |
| --- | --- | --- | --- | --- | --- |
| Fluorogold | Rabbit | Fluorochrome | Anti-Fluoro-Gold | 1:500 | Donkey Anti Rabbit Alexa Fluor 488 |
| NeuN | Guinea Pig | Millipore | ABN91 | 1:500 | Goat Anti-Guinea Pig Alexa Fluor 555 |
| ATP1A3 (α3) | Mouse | Abcam | ab2826 | 1:500 | Donkey Anti-Mouse Alexa Fluor 647 |
| ChAT | Goat | Millipore | Ab1-144P | 1:100 | Donkey Anti Goat CF488 |
| NeuN | Chicken | Millipore |  | 1:500 | Donkey anti Chicken Dylight 405 |
| Err3 | Mouse | Perseus Proteomics | PP-H6812-00 | 1:500 | Donkey anti Mouse Alexa 647 |
| Ankyrin G (Kizhatil *et al.*, 2007) | Rabbit | Gift- Paul Jenkins |  | 1:5000 | Donkey anti Rabbit 568 |

For AIS morphology, free-floating sections were first blocked in 5% donkey serum in PBS for 4 hours, followed by overnight incubation with a mixture of four primary antibodies (see Table 1) in PBS + 5% donkey serum. Sections were then sequentially incubated with corresponding secondary antibodies for 2 hours each (see Table 1). After final PBS washes, sections were mounted on Superfrost Plus slides and coverslipped using Dako Mounting Medium.

### Imaging and Analysis

For α3 expression, confocal images were acquired using an Andor BC43 Confocal Fluorescence Microscope with a 40x silicone objective lens (1.0 NA). Z-stacks were collected at 0.2 µm step size using four excitation wavelengths (405 nm, 488 nm, 560 nm, 640 nm). Image processing was performed in FIJI (Schindelin et al., 2012). Ten consecutive Z-planes were averaged into a single 2D projection. FG-positive motoneurons were identified by native FG fluorescence and/or FG immunolabeling. Somata were manually delineated to generate binarized masks, which were used to quantify fluorescence intensities for FG, NeuN, and α3 subunit. Membrane-specific α3 labeling was estimated by dilating the binary masks to capture perisomatic signal.

For AIS morphology, image stacks were obtained using a Zeiss LSM 700 confocal microscope with a 20x objective. Axon initial segments were identified by Ankyrin G immunolabeling and measured manually in three dimensions using Zen Black software (Zeiss).

### Research design and statistical analysis

Unpaired t-tests were performed to compare intrinsic properties and currents of putative gamma and slow alpha motoneurons. Appropriate and equivalent nonparametric tests (Mann-Whitney or Kruskal-Wallis) were conducted when data failed tests of normality or equal variance with Shapiro Wilk and Brown-Forsythe tests, respectively. Individual data points for all cells are presented in figures with mean ± SD. Statistical analyses were performed using Graph Pad Version 9.0 (Prism, San Diego, CA, USA). Results from all statistical tests are summarized in Table 2.

**Table 2:** Statistical Summary Table

|  | Source data | Condition | Property | No. of MNs | Statistical test | d.f. | Test statistic | P value | Hedge's g |
| --- | --- | --- | --- | --- | --- | --- | --- | --- | --- |
| a | 2B | gMN v. sMN | Rheobase | 18,21 | UPTT | 37 | 2.1 | 0.04 | 0.70 |
| b | 2C | gMN v. sMN | Capacitance | 18,21 | UPTT | 37 | 3.4 | 2.1e-3 | 0.90 |
| c | 2D | gMN v. sMN | Input Resist. | 18,21 | UPTT | 37 | 2.8 | 9.7e-3 | 0.76 |
| d | 2G | gMN v. sMN | Min. F.R. | 18,21 | UPTT | 37 | 2.8 | 9.4e-3 | 1.0 |
| e | 2H | gMN v. sMN | Max. F.R. | 18,21 | UPTT | 37 | 7.5 | 5.4e-8 | 2.5 |
| f | 2I | gMN v. sMN | F/I Gain | 18,21 | UPTT | 37 | 3.2 | 3.9e-3 | 1.15 |
| g | 3B | gMN v. sMN | Delta I | 12,15 | MWU |  | 62 | 0.2 | 0.59 |
| h | 3G | gMN v. sMN | PIC Amplitude | 6,7 | UPTT | 11 | 2.4 | 0.03 | 1.34 |
| i | text | gMN v. sMN | PIC Density | 6,7 | UPTT | 11 | 0.4 | 0.7 | 0.24 |
| j | 3H | gMN v. sMN | PIC on Volt. | 6,7 | UPTT | 11 | 2.4 | 0.03 | 1.42 |
| k | text | gMN v. sMN | RMP | 18,21 | UPTT | 37 | 2.4 | 0.02 | 0.76 |
| l | 4A2 | gMN v. sMN | Spike RT | 13,15 | MWU |  | 24 | 3.4e-4 | 1.13 |
| m | 4A3 | gMN v. sMN | Spike HW | 13,15 | UPTT | 26 | 3.2 | 3.2e-3 | 1.26 |
| n | 4B2 | gMN v. sMN | mAHP Amp. | 13,15 | UPTT | 26 | 2.3 | 2.8e-2 | 0.91 |
| o | 4B3 | gMN v. sMN | mAHP HW | 13,15 | UPTT | 26 | 0.2 | 0.8 | 0.01 |
| p | 5B | gMN vs. aMN | Soma size | 24,48 | UPTT | 70 | 10.2 | 2.0e-15 | 2.56 |
| q | 5C | gMN vs. aMN | AIS distance | 24,48 | UPTT | 70 | 4.3 | 6.6e-5 | 1.06 |
| r | 5D | gMN vs. aMN | AIS length | 24,48 | UPTT | 70 | 0.1 | 0.9 | 0 |
| s | 5E | gMN vs. aMN | Prox. Diameter | 24,48 | UPTT | 67 | 14.5 | 1.0e-15 | 3.55 |
| t | 5F | gMN vs. aMN | Dist. diameter | 24,48 | UPTT | 70 | 7.2 | 6.5e-10 | 1.77 |
| u | Text | gMN vs. aMN | Soma size | 6,6 | Wilcox. |  | -21.0 | 0.03 | 8.72 |
| v | Text | gMN vs. aMN | Prox. Diameter | 6,6 | Wilcox. |  | -21.0 | 0.03 | 7.12 |
| w | Text | gMN vs. aMN | Dist. diameter | 6,6 | Wilcox. |  | -21.0 | 0.03 | 3.31 |
| x | 6C2 | BL, Ouabain | usAHP Area | 6 | PTT | 5 | 5 | 0.03 | 1.62 |
| y | 6C3 | BL, Ouabain | usAHP Dur. | 6 | Wilcox. |  | -21 | 0.03 | 1.75 |
| z | 6C4 | BL, Ouabain | usAHP Amp. | 6 | Wilcox. |  | 9 | 0.4 | 0.27 |
| aa | 6E2 | gMN v. sMN | Area | 12,12 | MWU |  | 10 | 1.0e-4 | 1.94 |
| bb | 6E3 | gMN v. sMN | Duration | 12,12 | UPTT | 22 | 4.1 | 5.1e-4 | 1.66 |
| cc | 6E4 | gMN v. sMN | Amplitude | 12,12 | UPTT | 22 | 9 | 2.4e-3 | 1.40 |
| dd | 6F2 | gMN v. sMN | Ih Amplitude | 9,12 | UPTT | 19 | 0.5 | 0.6 | 0.23 |
| ee | 6F3 | gMN v. sMN | Ih Density | 9,12 | UPTT | 19 | 0.5 | 0.7 | 0.20 |
| ff | 7C2 | gMN v. sMN | A3 intensity | 25,35 | MWU |  | 376 | 0.4 |  |

**Table 3:** Intrinsic properties of putative gamma motoneurons and putative slow alpha motoneurons.

| Property | Putative gMN | Putative sMN | Effect Size | P value |
| --- | --- | --- | --- | --- |
| Rheobase (pA) | 122±98 | 229±187 | 0.70 | 0.04 |
| Capacitance (pF) | 191±71 | 276±111 | 0.90 | 2.1e-3 |
| Input Resistance (MΩ) | 149±129 | 75±57 | 0.76 | 9.7e-3 |
| Minimum Firing Rate (Hz) | 8.3±2.7 | 5.7±2.3 | 1.0 | 9.4e-3 |
| Maximum Firing Rate (Hz) | 115±33 | 51±15 | 2.5 | 5.4e-8 |
| F/I Gain (Hz/pA) | 0.16±0.1 | 0.06±0.06 | 1.15 | 3.9e-3 |
| Delta I (pA) | 67±114 | -5.7±130 | 0.59 | 0.2 |
| PIC Amplitude (pA) | -180±99 | -376±174 | 1.34 | 0.03 |
| PIC Density (pA/pF) | -1.0±0.6 | -1.1±0.5 | 0.24 | 0.7 |
| PIC Onset Voltage (mV) | -61±3.4 | -55±4.7 | 1.42 | 0.03 |
| Resting Membrane Potential (mV) | -59±6.4 | -65±7.5 | 0.76 | 0.02 |
| Spike Threshold (mV) | -44±3.8 | -44±4.7 | 0 | 0.9 |
| Spike Rise Time (ms) | 0.4±0.06 | 0.6±0.2 | 1.13 | 3.4e-4 |
| Spike Half Width (ms) | 0.4±0.07 | 0.6±0.1 | 1.26 | 3.2e-3 |
| mAHP Amplitude (mV) | -5.8±2.1 | -8.8±4.0 | 0.91 | 2.8e-2 |
| mAHP Half Width (ms) | 66±35 | 68±32 | 0.01 | 0.8 |
| Post-Discharge Area (mV*ms) | -78909±54296 | -1032±16726 | 1.94 | 1.0e-4 |
| Post-Discharge Duration (s) | 25.3±12.3 | 7.9±8.2 | 1.66 | 5.1e-4 |
| Post-Discharge Amplitude (mV) | -9.1±4.2 | -1.6±6.3 | 1.40 | 2.4e-3 |
| Max I<sub>h</sub> Amplitude (pA) | -333±299 | -390±180 | 0.23 | 0.6 |
| I<sub>h</sub> Density (pA/pF) | -1.5±1.0 | -1.3±0.6 | 0.20 | 0.7 |

## Contributions

Study Conception and Design: SAS, GBM; Data acquisition and analysis: SAS, SJN, DBJ, MJB, FLS; Preparation of Figures: SAS; First draft of manuscript: SAS, GBM; Funding and supervision: SAS, CFM, GBM. All authors approved the final version of the manuscript.

## Acknowledgements

The authors reserve the right to apply a Creative Commons Attribution (CC BY) licence to any Author Accepted manuscript version arising from this submission. The research data supporting this publication will be made freely available using an open access data repository following acceptance to a peer-reviewed journal. The Core Facility for Integrated Microscopy, Faculty of Health and Medical Sciences, University of Copenhagen for imaging. We also thank Paul Jenkins (The university of Michigan) and Peter Mohler (The Ohio State University) for the gift of the Ankyrin G antibody.

## Data Availability Statement

Data will be made available through the Open Science Framework providing a DOI to this source for this paper once accepted to a peer reviewed journal. A data availability statement can be found in the Acknowledgements statement and also in the last sentence of the Methods.

## References

1. Akkuratov EE, Sorrell F, Picton LD, Sousa VC, Paucar M, Jans D, Svensson L-B, Lindskog M, Fritz N, Liebmann T, Sillar KT, Rosewich H, Svenningsson P, Brismar H, Miles GB & Aperia A (2025). ATP1A3 dysfunction causes motor hyperexcitability and afterhyperpolarization loss in a dystonia model. Brain 148, 1099–1105.

2. Alkaslasi MR, Piccus ZE, Hareendran S, Silberberg H, Chen L, Zhang Y, Petros TJ & Le Pichon CE (2021). Single nucleus RNA-sequencing defines unexpected diversity of cholinergic neuron types in the adult mouse spinal cord. Nat Commun 12, 2471.

3. Altman J & Sudarshan K (1975). Postnatal development of locomotion in the laboratory rat. Anim Behav 23, 896–920.

4. Ashrafi S, Lalancette-Hébert M, Friese A, Sigrist M, Arber S, Shneider NA & Kaltschmidt JA (2012). Wnt7A identifies embryonic γ-motor neurons and reveals early postnatal dependence of γ-motor neurons on a muscle spindle-derived signal. J Neurosci 32, 8725–8731.

5. Bakels R & Kernell D (1993). Average but not continuous speed match between motoneurons and muscle units of rat tibialis anterior. J Neurophysiol 70, 1300–1306.

6. Bellingham MC (2013). Pre- and postsynaptic mechanisms underlying inhibition of hypoglossal motor neuron excitability by riluzole. J Neurophysiol 110, 1047–1061.

7. Bennett DJ, Li Y & Siu M (2001). Plateau potentials in sacrocaudal motoneurons of chronic spinal rats, recorded in vitro. J Neurophysiol 86, 1955–1971.

8. Bhumbra GS & Beato M (2018). Recurrent excitation between motoneurones propagates across segments and is purely glutamatergic. PLoS Biol 16, e2003586.

9. Blum JA, Klemm S, Shadrach JL, Guttenplan KA, Nakayama L, Kathiria A, Hoang PT, Gautier O, Kaltschmidt JA, Greenleaf WJ & Gitler AD (2021). Single-cell transcriptomic analysis of the adult mouse spinal cord reveals molecular diversity of autonomic and skeletal motor neurons. Nat Neurosci 24, 572–583.

10. Bos R, Drouillas B, Bouhadfane M, Pecchi E, Trouplin V, Korogod SM & Brocard F (2021). Trpm5 channels encode bistability of spinal motoneurons and ensure motor control of hindlimbs in mice. Nat Commun 12, 6815.

11. Bos R, Harris-Warrick RM, Brocard C, Demianenko LE, Manuel M, Zytnicki D, Korogod SM & Brocard F (2018). Kv1.2 Channels Promote Nonlinear Spiking Motoneurons for Powering Up Locomotion. Cell Rep 22, 3315–3327.

12. Bouhadfane M, Tazerart S, Moqrich A, Vinay L & Brocard F (2013). Sodium-mediated plateau potentials in lumbar motoneurons of neonatal rats. J Neurosci 33, 15626–15641.

13. Brocard C, Plantier V, Boulenguez P, Liabeuf S, Bouhadfane M, Viallat-Lieutaud A, Vinay L & Brocard F (2016). Cleavage of Na(+) channels by calpain increases persistent Na(+) current and promotes spasticity after spinal cord injury. Nat Med 22, 404–411.

14. Carlin KP, Jiang Z & Brownstone RM (2000a). Characterization of calcium currents in functionally mature mouse spinal motoneurons. Eur J Neurosci 12, 1624–1634.

15. Carlin KP, Jones KE, Jiang Z, Jordan LM & Brownstone RM (2000b). Dendritic L-type calcium currents in mouse spinal motoneurons: implications for bistability. Eur J Neurosci 12, 1635– 1646.

16. Deardorff AS, Romer SH, Deng Z, Bullinger KL, Nardelli P, Cope TC & Fyffe REW (2013). Expression of postsynaptic Ca2+-activated K+ (SK) channels at C-bouton synapses in mammalian lumbar -motoneurons. J Physiol 591, 875–897.

17. Djukic S, Zhao Z, Jørgensen LMH, Bak AN, Jensen DB & Meehan CF (2025). TDP-43 pathology is sufficient to drive axon initial segment plasticity and hyperexcitability of spinal motoneurones in vivo in the TDP43-ΔNLS model of Amyotrophic Lateral Sclerosis. Acta Neuropathol Commun 13, 42.

18. Drouillas B, Brocard C, Zanella S, Bos R & Brocard F (2023). Persistent Nav1.1 and Nav1.6 currents drive spinal locomotor functions through nonlinear dynamics. Cell Rep 42, 113085.

19. Durand J, Filipchuk A, Pambo-Pambo A, Amendola J, Borisovna Kulagina I & Guéritaud J-P (2015). Developing electrical properties of postnatal mouse lumbar motoneurons. Front Cell Neurosci 9, 349.

20. Eccles JC (1957). Some aspects of Sherrington’s contribution to neurophysiology. Notes Rec R Soc Lond 12, 216–225.

21. Eccles JC, Eccles RM & Lundberg A (1957). Durations of after-hyperpolarization of motoneurones supplying fast and slow muscles. Nature 179, 866–868.

22. Edwards IJ, Bruce G, Lawrenson C, Howe L, Clapcote SJ, Deuchars SA & Deuchars J (2013). Na+/K+ ATPase α1 and α3 isoforms are differentially expressed in α- and γ-motoneurons. J Neurosci 33, 9913–9919.

23. Elbasiouny SM, Bennett DJ & Mushahwar VK (2005). Simulation of dendritic CaV1.3 channels in cat lumbar motoneurons: spatial distribution. J Neurophysiol 94, 3961–3974.

24. Elbasiouny SM, Bennett DJ & Mushahwar VK (2006). Simulation of Ca2+ persistent inward currents in spinal motoneurones: mode of activation and integration of synaptic inputs. J Physiol 570, 355–374.

25. Eleftheriadis PE, Pothakos K, Sharples SA, Apostolou PE, Mina M, Tetringa E, Tsape E, Miles GB & Zagoraiou L (2023). Peptidergic modulation of motor neuron output via CART signaling at C bouton synapses. Proc Natl Acad Sci U S A 120, e2300348120.

26. Ellaway PH, Taylor A & Durbaba R (2015). Muscle spindle and fusimotor activity in locomotion. J Anat 227, 157–166.

27. Enjin A, Leão KE, Mikulovic S, Le Merre P, Tourtellotte WG & Kullander K (2012). Sensorimotor function is modulated by the serotonin receptor 1d, a novel marker for gamma motor neurons. Mol Cell Neurosci 49, 322–332.

28. Fékété A, Ankri N, Brette R & Debanne D (2021). Neural excitability increases with axonal resistance between soma and axon initial segment. Proc Natl Acad Sci U S A; DOI: 10.1073/pnas.2102217118.

29. Friese A, Kaltschmidt JA, Ladle DR, Sigrist M, Jessell TM & Arber S (2009). Gamma and alpha motor neurons distinguished by expression of transcription factor Err3. Proc Natl Acad Sci U S A 106, 13588–13593.

30. Gardiner PF & Kernell D (1990). The “fastness” of rat motoneurones: time-course of afterhyperpolarization in relation to axonal conduction velocity and muscle unit contractile speed. Pflugers Arch 415, 762–766.

31. Goethals S & Brette R (2020). Theoretical relation between axon initial segment geometry and excitability. Elife; DOI: 10.7554/eLife.53432.

32. Granit R (1975). The functional role of the muscle spindles--facts and hypotheses. Brain 98, 531–556.

33. Gulledge AT & Bravo JJ (2016). Neuron Morphology Influences Axon Initial Segment Plasticity. eNeuro; DOI: 10.1523/ENEURO.0085-15.2016.

34. Gustafsson B & Lipski J (1979). Do gamma-motoneurons lack a long-lasting afterhyperpolarization? Brain Res 172, 349–353.

35. Harrison PJ (1983). The relationship between the distribution of motor unit mechanical properties and the forces due to recruitment and to rate coding for the generation of muscle force. Brain Res 264, 311–315.

36. Harris-Warrick RM, Pecchi E, Drouillas B, Brocard F & Bos R (2024). Effect of size on expression of bistability in mouse spinal motoneurons. J Neurophysiol 131, 577–588.

37. Heckmann CJ, Gorassini MA & Bennett DJ (2005). Persistent inward currents in motoneuron dendrites: implications for motor output. Muscle Nerve 31, 135–156.

38. Henneman E (1957). Relation between size of neurons and their susceptibility to discharge. Science 126, 1345–1347.

39. Hochman S (2011). Long-term patch recordings from adult spinal neurons herald new era of opportunity. J Neurophysiol 106, 2794–2795.

40. Jensen DB, Klingenberg S, Dimintiyanova KP, Wienecke J & Meehan CF (2020). Intramuscular Botulinum toxin A injections induce central changes to axon initial segments and cholinergic boutons on spinal motoneurones in rats. Sci Rep 10, 893.

41. Jiang Z, Rempel J, Li J, Sawchuk MA, Carlin KP & Brownstone RM (1999). Development of L-type calcium channels and a nifedipine-sensitive motor activity in the postnatal mouse spinal cord. Eur J Neurosci 11, 3481–3487.

42. Jørgensen HS, Jensen DB, Dimintiyanova KP, Bonnevie VS, Hedegaard A, Lehnhoff J, Moldovan M, Grondahl L & Meehan CF (2021). Increased Axon Initial Segment Length Results in Increased Na Currents in Spinal Motoneurones at Symptom Onset in the G127X SOD1 Mouse Model of Amyotrophic Lateral Sclerosis. Neuroscience 468, 247–264.

43. Kang Y, Saito M & Toyoda H (2024). Molecular, Morphological and Electrophysiological Differences between Alpha and Gamma Motoneurons with Special Reference to the Trigeminal Motor Nucleus of Rat. Int J Mol Sci; DOI: 10.3390/ijms25105266.

44. Karekal A, Mandawe R, Chun C, Byri SK, Cheline D, Ortiz S, Hochman S & Wilkinson KA (2025). Optogenetic methods to stimulate gamma motor neuron axons ex vivo. Exp Physiol; DOI: 10.1113/EP092359.

45. Kemm RE & Westbury DR (1978). Some properties of spinal gamma-motoneurones in the cat, determined by micro-electrode recording. J Physiol 282, 59–71.

46. Kernell D (1965). The limits of firing frequency in cat lumbosacral motoneurones possessing different time course of afterhyperpolarization. Acta Physiol Scand 65, 87–100.

47. Kernell D, Bakels R & Copray JC (1999). Discharge properties of motoneurones: how are they matched to the properties and use of their muscle units? J Physiol Paris 93, 87–96.

48. Khan MN, Cherukuri P, Negro F, Rajput A, Fabrowski P, Bansal V, Lancelin C, Lee T-I, Bian Y, Mayer WP, Akay T, Müller D, Bonn S, Farina D & Marquardt T (2022). ERR2 and ERR3 promote the development of gamma motor neuron functional properties required for proprioceptive movement control. PLoS Biol 20, e3001923.

49. Kissane RWP, Ghaffari-Rafi A, Tickle PG, Chakrabarty S, Egginton S, Brownstone RM & Smith CC (2022). C-bouton components on rat extensor digitorum longus motoneurons are resistant to chronic functional overload. J Anat 241, 1157–1168.

50. Kizhatil K, Davis JQ, Davis L, Hoffman J, Hogan BLM & Bennett V (2007). Ankyrin-G is a molecular partner of E-cadherin in epithelial cells and early embryos. J Biol Chem 282, 26552– 26561.

51. Koronfel LM, Kanning KC, Alcos A, Henderson CE & Brownstone RM (2021). Elimination of glutamatergic transmission from Hb9 interneurons does not impact treadmill locomotion. Sci Rep 11, 16008.

52. Kussick E et al. (2025). Enhancer AAVs for targeting spinal motor neurons and descending motor pathways in rodents and macaque. Cell Rep115730.

53. Lee RH & Heckman CJ (1998). Bistability in spinal motoneurons in vivo: systematic variations in persistent inward currents. J Neurophysiol 80, 583–593.

54. Leroy F, Lamotte d’Incamps B, Imhoff-Manuel RD & Zytnicki D (2014). Early intrinsic hyperexcitability does not contribute to motoneuron degeneration in amyotrophic lateral sclerosis. Elife; DOI: 10.7554/eLife.04046.

55. Leroy F, Lamotte d’Incamps B & Zytnicki D (2015). Potassium currents dynamically set the recruitment and firing properties of F-type motoneurons in neonatal mice. J Neurophysiol 114, 1963–1973.

56. Li Y & Bennett DJ (2003). Persistent sodium and calcium currents cause plateau potentials in motoneurons of chronic spinal rats. J Neurophysiol 90, 857–869.

57. Li Y, Gorassini MA & Bennett DJ (2004). Role of persistent sodium and calcium currents in motoneuron firing and spasticity in chronic spinal rats. J Neurophysiol 91, 767–783.

58. Manuel M & Zytnicki D (2019). Molecular and electrophysiological properties of mouse motoneuron and motor unit subtypes. Curr Opin Physiol 8, 23–29.

59. Miles GB, Dai Y & Brownstone RM (2005). Mechanisms underlying the early phase of spike frequency adaptation in mouse spinal motoneurones. J Physiol 566, 519–532.

60. Miles GB, Hartley R, Todd AJ & Brownstone RM (2007). Spinal cholinergic interneurons regulate the excitability of motoneurons during locomotion. Proc Natl Acad Sci U S A 104, 2448–2453.

61. Mitra P & Brownstone RM (2012). An in vitro spinal cord slice preparation for recording from lumbar motoneurons of the adult mouse. J Neurophysiol 107, 728–741.

62. Morita-Isogai Y, Sato H, Saito M, Kuramoto E, Yin DX, Kaneko T, Yamashiro T, Takada K, Oh SB, Toyoda H & Kang Y (2017). A distinct functional distribution of α and γ motoneurons in the rat trigeminal motor nucleus. Brain Struct Funct 222, 3231–3239.

63. Nakanishi ST & Whelan PJ (2010). Diversification of intrinsic motoneuron electrical properties during normal development and botulinum toxin-induced muscle paralysis in early postnatal mice. J Neurophysiol 103, 2833–2845.

64. Nascimento F, Broadhead MJ, Tetringa E, Tsape E, Zagoraiou L & Miles GB (2020). Synaptic mechanisms underlying modulation of locomotor-related motoneuron output by premotor cholinergic interneurons. Elife; DOI: 10.7554/eLife.54170.

65. Nascimento F, Özyurt MG, Halablab K, Bhumbra GS, Caron G, Bączyk M, Zytnicki D, Manuel M, Roselli F, Brownstone R & Beato M (2024). Spinal microcircuits go through multiphasic homeostatic compensations in a mouse model of motoneuron degeneration. Cell Rep 43, 115046.

66. Nishimura K, Ohta M, Saito M, Morita-Isogai Y, Sato H, Kuramoto E, Yin DX, Maeda Y, Kaneko T, Yamashiro T, Takada K, Oh SB, Toyoda H & Kang Y (2018). Electrophysiological and Morphological Properties of α and γ Motoneurons in the Rat Trigeminal Motor Nucleus. Front Cell Neurosci 12, 9.

67. Niyo G, Almofeez LI, Erwin A & Valero-Cuevas FJ (2024). A computational study of how an α- to γ-motoneurone collateral can mitigate velocity-dependent stretch reflexes during voluntary movement. Proc Natl Acad Sci U S A 121, e2321659121.

68. Özyurt MG, Ojeda-Alonso J, Beato M & Nascimento F (2022). In vitro longitudinal lumbar spinal cord preparations to study sensory and recurrent motor microcircuits of juvenile mice. J Neurophysiol 128, 711–726.

69. Patel T, Hammelman J, Aziz S, Jang S, Closser M, Michaels TL, Blum JA, Gifford DK & Wichterle H (2022). Transcriptional dynamics of murine motor neuron maturation in vivo and in vitro. Nat Commun 13, 5427.

70. Picton LD, Nascimento F, Broadhead MJ, Sillar KT & Miles GB (2017). Sodium Pumps Mediate Activity-Dependent Changes in Mammalian Motor Networks. J Neurosci 37, 906–921.

71. Picton LD, Sillar KT & Zhang H-Y (2018). Control of Xenopus Tadpole Locomotion via Selective Expression of Ih in Excitatory Interneurons. Curr Biol 28, 3911–3923.e2.

72. Pulver SR & Griffith LC (2010). Spike integration and cellular memory in a rhythmic network from Na+/K+ pump current dynamics. Nat Neurosci 13, 53–59.

73. Quinlan KA, Schuster JE, Fu R, Siddique T & Heckman CJ (2011). Altered postnatal maturation of electrical properties in spinal motoneurons in a mouse model of amyotrophic lateral sclerosis. J Physiol 589, 2245–2260.

74. Reedich EJ, Genry LT, Steele PR, Mena Avila E, Dowaliby L, Drobyshevsky A, Manuel M & Quinlan KA (2023). Spinal motoneurons respond aberrantly to serotonin in a rabbit model of cerebral palsy. J Physiol 601, 4271–4289.

75. Rotterman TM, Carrasco DI, Housley SN, Nardelli P, Powers RK & Cope TC (2021). Axon initial segment geometry in relation to motoneuron excitability. PLoS One 16, e0259918.

76. Schwindt P & Crill WE (1977). A persistent negative resistance in cat lumbar motoneurons. Brain Res 120, 173–178.

77. Sharples SA, Broadhead MJ, Gray JA & Miles GB (2023). M-type potassium currents differentially affect activation of motoneuron subtypes and tune recruitment gain. J Physiol 601, 5751–5775.

78. Sharples SA & Miles GB (2021). Maturation of persistent and hyperpolarization-activated inward currents shapes the differential activation of motoneuron subtypes during postnatal development. Elife; DOI: 10.7554/eLife.71385.

79. Shneider NA, Brown MN, Smith CA, Pickel J & Alvarez FJ (2009). Gamma motor neurons express distinct genetic markers at birth and require muscle spindle-derived GDNF for postnatal survival. Neural Dev 4, 42.

80. Smith CC & Brownstone RM (2020). Spinal motoneuron firing properties mature from rostral to caudal during postnatal development of the mouse. J Physiol 598, 5467–5485.

81. Steele PR, Cavarsan CF, Dowaliby L, Westefeld M, Katenka N, Drobyshevsky A, Gorassini MA & Quinlan KA (2020). Altered Motoneuron Properties Contribute to Motor Deficits in a Rabbit Hypoxia-Ischemia Model of Cerebral Palsy. Front Cell Neurosci 14, 69.

82. Verneuil J, Brocard C, Trouplin V, Villard L, Peyronnet-Roux J & Brocard F (2020). The M-current works in tandem with the persistent sodium current to set the speed of locomotion. PLoS Biol 18, e3000738.

83. Westbury DR (1981). Electrophysiological characteristics of spinal gamma motoneurons in the cat. In Muscle Receptors and Movement, pp. 87–96. Palgrave Macmillan UK, London.

84. Wilkinson KA (2021). Methodological advances for studying gamma motor neurons. Curr Opin Physiol 19, 135–140.

85. Zhang H-Y, Picton L, Li W-C & Sillar KT (2015). Mechanisms underlying the activity-dependent regulation of locomotor network performance by the Na+ pump. Sci Rep 5, 16188.

