## Supplementary Figure 1 for "Intrinsic mechanisms contributing to the biophysical signature of mouse gamma motoneurons"

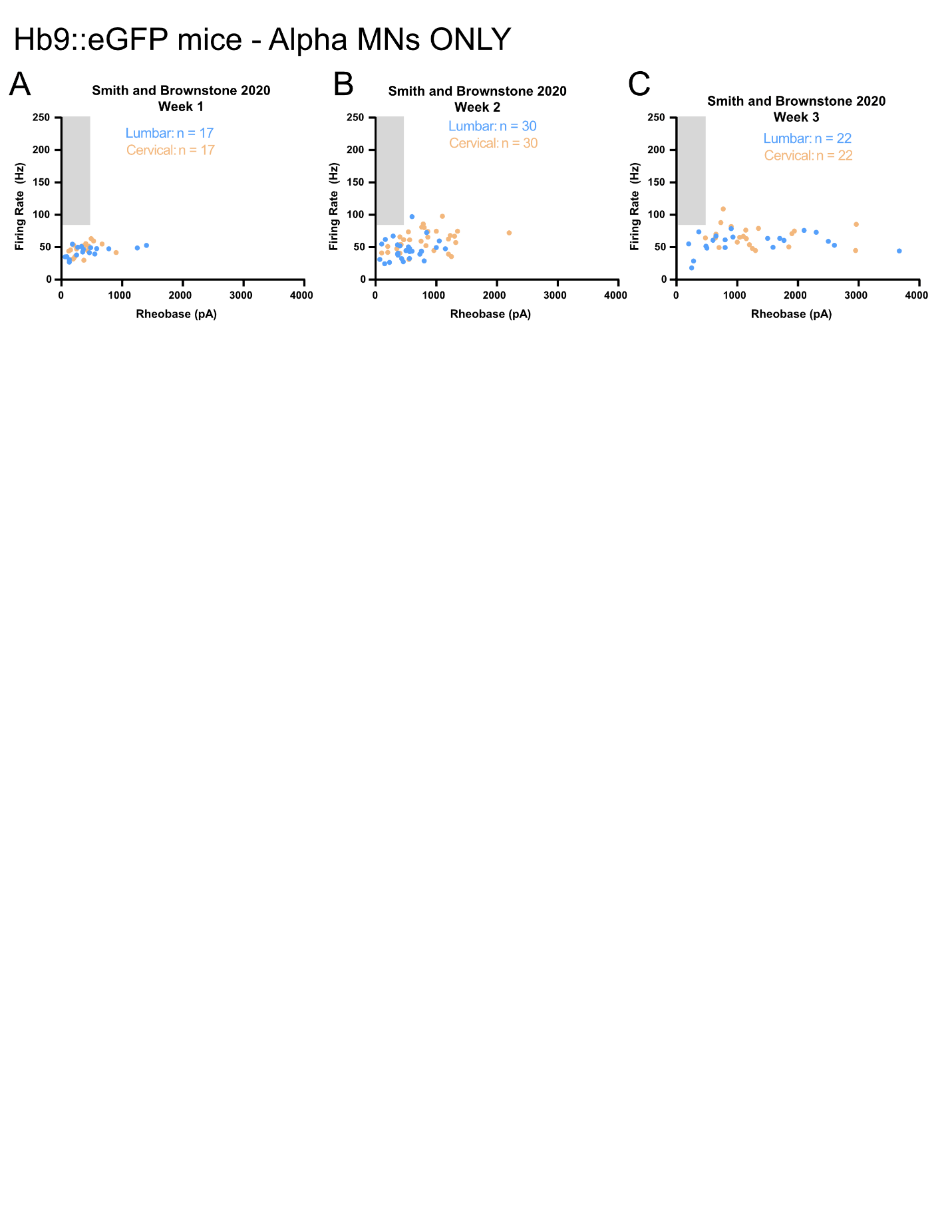

**Supplementary Figure 1:** A-C) Maximum firing rate assessed during depolarizing current steps and plotted as a function of rheobase for Hb9::GFP+ motoneurons in lumbar (blue) and cervical (orange) spinal segments during postnatal weeks 1-3 studied by Smith and Brownstone (2020).
